# Investigate Channel Rectifications and Neural Dynamics by An Electrodiffusive Gauss-Nernst-Planck Approach

**DOI:** 10.1101/2025.02.18.638790

**Authors:** Zichao Liu, Yinyun Li

**Affiliations:** School of Systems Science, Beijing Normal University, Xinjiekou Wai Street 19. Beijing, 100875, China; Computational Neuroscience Unit, Okinawa Institute of Science and Technology, 1919-1, Tancha, Onna-son, Kunigami-gun, Okinawa, Japan

## Abstract

Electrodiffusion plays a crucial role in modulating ion channel conductivity and neural firing dynamics within the nervous system. However, its influence on neural function remains underexplored. In this work, we introduce a novel Gauss-Nernst-Planck (GNP) approach to investigate how electrodiffusive dynamics influence ion channel rectification and neural activity. For the first time, we demonstrate that membrane conductivity is determined by both ion permeability and the harmonic mean of intramembrane ion concentration, bridging the gap between the permeability-based Goldman-Hodgkin-Katz (GHK) model and conductance-based models. We characterize the rectification properties of ***GABA***_***A***_, ***AMPA***, and leaky channels by estimating their single-channel permeabilities and conductance. By integrating these rectifying channels into neurodynamic models, our GNP neurodynamic model reveals how electrodiffusive dynamics fundamentally shape neural firing by modulating membrane conductance and the interplay between passive and active ion transport-mechanisms that are often overlooked in conventional conductance-based neurodynamic models. This study provides a fundamental mechanistic understanding of electrodiffusion regulation in neural activity and establishes a robust framework for future research in neurophysiology.

## Introduction

The transmembrane movement of ions alters the electric potential across the neural membrane, forming the basis of various physiological and pathological neural behaviors [1, 2]. Unlike the movement of electrons in conductors, which is solely driven by electric potential differences, ion fluxes in the neural system are influenced by both electric potential and concentration gradients, collectively characterized as “electrodiffusive movement” [3, 4]. The unique properties of transmembrane electrodiffusion are captured by the well-known Goldman-Hodgkin-Katz (GHK) current equation, which explains the nonlinear I-V relationships observed in various ion channels, a phenomenon known as Goldman rectification [5,6]. However, the extent to which ion electrodiffusion influences neural activity and excitability remains insufficiently explored [7].

A major limitation in studies of electrodiffusion in the neural system is that, while Goldman rectification has been extensively examined across various ion channels [8, 9], electrodiffusive dynamics are often overlooked at the membrane or whole-neuron level. In computational models, transmembrane electrodiffusive ion fluxes are typically simplified as electrical currents in circuit-based representations [10–12]. This simplification disregards the underlying electrodiffusive processes, even though neural activity fundamentally relies on ion channels exhibiting significant rectification properties. Given that ion channel rectification plays a critical role in shaping neural function [13], its omission in traditional conductance-based models may lead to inaccurate simulations and misinterpretations of neural electrophysiology. This issue becomes particularly concerning in pathological conditions such as epileptic seizures and cortical spreading depression, where ion concentration gradients undergo dramatic shifts [14, 15]. These shifts reshape transmembrane electrodiffusion and alter channel rectification properties, yet their impact on pathological neural dynamics remains largely unexplored. Addressing these challenges requires a deeper investigation into transmembrane electrodiffusion to refine our understanding of neural function.

Incorporating electrodiffusive dynamics into neurodynamic models, however, remains a significant challenge. While many ion channels exhibit rectification properties, others, such as α-Amino-3-hydroxy-5-methyl-4-isoxazolepropionic acid (***AMPA***) and ***NALCN*** channels, display linear I-V relationships [16, 17]. The alignment of these linear I-V curves with circuit equations raises questions about whether electrodiffusive dynamics are necessary for modeling currents through such channels. Additionally, certain ion channels, such as Gamma-Aminobutyric Acid Type A (***GABA***_***A***_) receptors, exhibit strong outward rectification under physiological conditions, yet quantifying their single-channel conductance remains difficult [18–20]. Some studies estimate ***GABA***_***A***_ channel conductance by measuring the slope of I-V curves [18], but the validity of this approach is debated, and it is not easily applicable in constructing whole-cell neurodynamic models.

Many studies have investigated electrodiffusion in the nervous system by solving the Poisson-Nernst-Planck (PNP) equations [21, 22], providing high-resolution simulations of ion transport across ion channels [23–25], between the soma and dendrites [26, 27], and within extracellular spaces [28–31]. However, solving PNP equations requires substantial computational resources, making them impractical for whole-neuron dynamic modeling. As a result, a practical neurodynamic model that effectively incorporates electrodiffusive dynamics remains lacking.

In this paper, we introduce a novel Gauss-Nernst-Planck (GNP) framework that models the electrodiffusive transmembrane ion dynamics in regulating channel properties and neural dynamics. Our GNP approach reveals the intrinsic quantitative relation between membrane conductivity and permeability, effectively bridging the gap between the permeability-based GHK current equation and the conductance-based circuit models. Using this method, we estimate the permeabilities and conductance of ***GABA***_***A***_ and ***AMPA*** channels for their respective permeant ions. We elucidate the mechanisms behind the outward rectification of ***GABA***_***A***_ channels [18–20], the linear I-V relationship of GluR2-containing ***AMPA*** channels [16, 32], and the inward rectification of GluR2-lacking ***AMPA*** channels [33, 34]. Additionally, we developed a neurodynamic model incorporating rectifying ion channels to explore how electrodiffusive dynamics fundamentally modulate neural firing activity through the interplay between passive and active ion transport. We conclude our main results and discuss the potential application of our model in the Discussion.

## Results

### Determine the Membrane Conductivity Through Electrodiffusive Dynamics

We introduce an innovative approach to integrating intramembrane electrodiffusive dynamics into neural system modeling.

By assuming overall electrical neutrality in the neural system (Fig. 1a, 1b), we leverage Gauss’ s Law to replace the Poisson equation in the Poisson-Nernst-Planck (PNP) framework. This key simplification not only reduces the complexity of the PNP problem but also establishes a crucial link between electrodiffusive- and conductance-based models (see Supporting Information for details).

**Figure 1.**
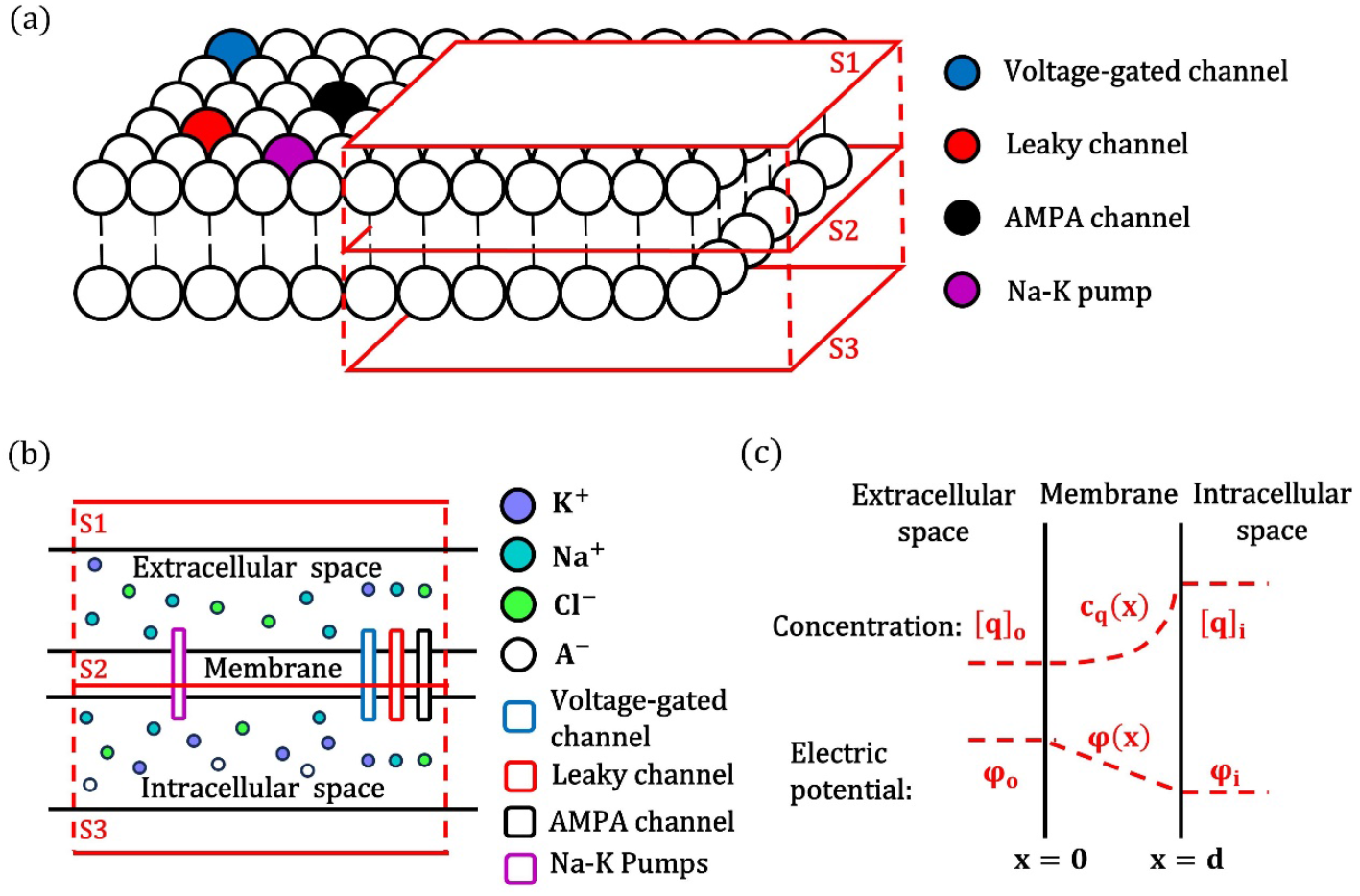
Schematic illustration of the Gauss-Nernst-Planck Approach. (a) Neuronal membrane with various channels and ***Na***^**+**^**/*K***^**+**^ **− *ATPase*** inserted into the parallel bilayer-lipid structure. Surfaces ***S*1**, ***S*2**, and ***S*3** are parallel, with ***S*2** located within the intramembrane space. (b) Illustration of space division for the neuron membrane. The intracellular and extracellular spaces contain different ion concentrations **(*K***^**+**^, ***Na***^**+**^, ***Cl***^**−**^), ions move through membrane by electrodiffusion via channels such as leaky channel, voltage-gated channel and also by active transport of ***Na***^**+**^**/*K***^**+**^**− *ATPase. A***^**−**^represents impermeable anions confined to the intracellular space. (c) The concentration profile of ion (***c***_***q***_**(*x*)**) and electric potential distribution (***φ*(*x*)**) in the intramembrane space. The ion concentration profile is nonlinear, while the electric potential profile is linear by Gauss’s law. Extracellular space: ***x* < 0**, intracellular space: ***x* > *d***. intramembrane space: ***x* ∈ [0, *d*]**.

Through this Gauss-Nernst-Planck framework, we demonstrate that membrane conductance (per unit area) can be expressed as a function of permeability and the intramembrane concentration profile (denoted as *c*_*q*_(*x*), see Fig. 1c) of the permeant ion. This relationship provides a fundamental link between ion electrodiffusion dynamics and membrane properties, as shown in Eq. 1:

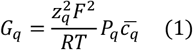

Where, *R* is the gas constant, *T* is temperature, and 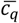 is the harmonic mean of intramembrane concentration profile *c*_*q*_(*x*). At steady state, *c*_*q*_(*x*) is determined by both ion concentrations and membrane potential (see Eq. 13). As shown in Fig. 2a-2d, variations in membrane potential result in changes in the concentration profiles of potassium, sodium, chloride, and calcium ions, which subsequently alter membrane conductance and induce Goldman rectification.

**Figure 2.**
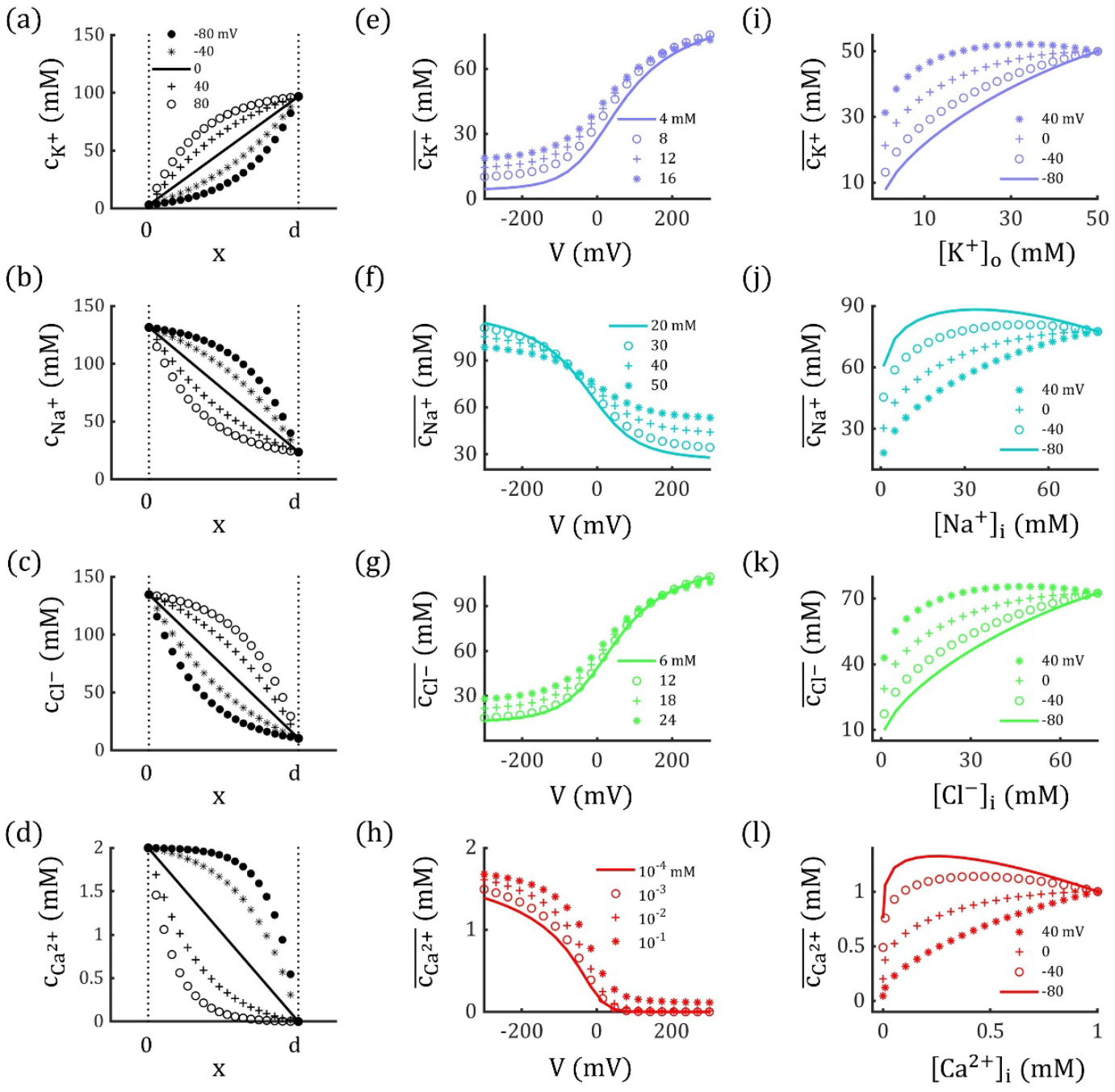
Ion Concentration Profiles *c*_*q*_(*x*) and Corresponding Harmonic Means 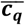 as Functions of Membrane Potential and Ion Concentrations. (a-d) Variation of concentration profiles for potassium, sodium, chloride, and calcium ions with different membrane potentials. (e-h) Harmonic mean of each ion concentration profile changes with membrane potential. As membrane potential increases, 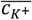 and 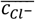 increase, resulting in outward rectification, whereas 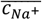 and 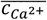 decrease, leading to inward rectification. (i-l) Harmonic means of each ion concentration profile typically increases as the concentration gradient **Δ*c***_***q***_ **=** |**[*q*]**_***i***_ **− [*q*]**_***o***_| decreases. The total ion concentration (**[*q*]**_***i***_ **+ [*q*]**_***o***_) is held constant.

Importantly, Eq. 1 bridges the conductance-based model and permeability-based electrodiffusive approach [35], which allows us to model neural firing activity by both circuit-like and GHK current equations through GNP approach, and investigate the effect of electrodiffusion on neural firing properties (see Methods).

To further illustrate how concentration profiles influence conductivity, we derived an analytical expression for 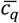 by integrating *c*_*q*_(*x*) (see Methods):

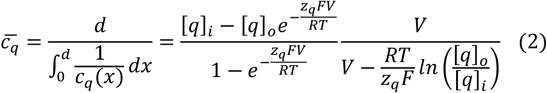

Eq. 2 highlights that both membrane potential and ion concentrations affect 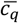, thereby altering conductance by Eq. 1. As shown in Fig. 2e–2h, the harmonic means of the concentration profiles for potassium and chloride ions 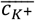 and 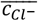 increase with membrane potentials, whereas 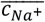 and 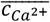 decrease with it. These shifts lead to higher conductance for inhibitory currents and lower conductance for excitatory currents during depolarization. This differential conductance produces outward rectification for potassium and chloride currents and inward rectification for sodium and calcium currents. The voltage dependent properties suggest dynamic alterations of membrane potential will also modify the channel conductance by altering 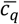, creating a feedback loop that is absent in traditional conductance-based neurodynamic models.

Moreover, as illustrated in Fig. 2i–2l, 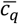 typically increases as the concentration gradient Δ*c*_*q*_ = |[*q*]_*i*_ − [*q*]_*o*_| decreases, which subsequently upregulates the associated conductance. The variation of 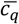 also diminishes with decreasing Δ*c*_*q*_, reducing rectification and eventually producing a linear I-V relation when Δ*c*_*q*_ = 0 *mM*. During physiologically normal neural activity, homeostatic mechanisms help stabilize Δ*c*_*q*_ [36], which may explain why the concentration-dependence of membrane conductance is often overlooked. However, under pathological conditions like depolarization blocks, Δ*c*_*q*_ can change significantly [36–38], which would be illustrated in Fig. 6.

### Characterizing Channel Rectification by GNP Approach

The GHK current equation has been widely applied to investigate the biophysical and electrical properties of ion channels, particularly because many channels exhibit Goldman rectification. However, without accurately characterizing the permeability and conductivity of channels, the application of their rectification properties in neurodynamic models remains challenging.

Some of the key unresolved questions include

1. Conductance of rectifying channel: Many channels, such as ***GABA***_***A***_ channels [18–20], display rectification, rendering their voltage-dependent conductance difficult to be accurately determined. While some studies derive the conductance by taking the slope of the I-V curve (the derivative of the GHK current equation with respect to voltage), this method is neither effective nor precise.
2. Single channel permeability: The single-channel conductance of ion channels is measured through their linear I-V curve, such as ***AMPA*** channels [16, 32]. However, as shown in Eq. 2, the conductivity is influenced by various factors, making it less informative to characterize the channel’s intrinsic biophysical properties than channel permeability. However, ion permeability data for channels are rarely measured.
3. Mechanisms of rectification: Channels like ***AMPA*** and ***NALCN*** typically exhibit a linear I-V relationship [16, 17, 32]. However, under certain conditions, such as ***AMPA*** channels lacks GluR2 subunit and ion concentrations are changed around ***NALCN*** channels, these channels could show rectification [17, 33, 34]. The underlying mechanism driving this shift remains unclear.
4. Specifying ionic currents: Many ion channels are permeable to multiple ion types. While total current is easily measurable, separating the ionic currents for each ion type is challenging. It is often assumed that the respective ionic currents are proportional to permeability ratios [16, 19], but this assumption is problematic due to the non-linear functions revealed by Eqs. 1, 2.

To address those challenges, we propose a practical methodology to determine the conductivity and permeability of ion channels based on available data. First, we determine how to achieve a linear I-V relationship by adjusting ion concentrations in individual ion channels (see Eq. 15 in Methods). Next, we calculate single-channel permeabilities using single-channel conductance (the slope of the linear I-V curve) and ion concentration data (see Eq. 16 in Methods). Finally, we quantify channel conductance for both specific permeant ion currents and total current (see Eqs. 18–25 in Methods). We will use this framework to address the issues mentioned above, by taking ***GABA***_***A***_ channel and ***AMPA*** channel as examples.

#### *GABA*_*A*_ Channel

To apply the GNP framework to analyze channel rectification properties, the required information includes permeability ratios, single-channel conductance, the pore diameter of the channel, and permeants’ concentrations in the intra- and extracellular spaces. All those data have been reported in previous experimental research.

For ***GABA***_***A***_ channel, both chloride and bicarbonate ions as permeants with valence of −1. Experimental studies suggest that the permeability ratio 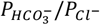 is approximately 0.2, and the pore diameter of the channel is around 5 Å [19, 20]. The concentrations of 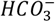 ions are set at 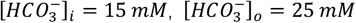[36].

In experiments, to measure the single-channel conductance of ***GABA***_***A***_ channel, both [*Cl*^−^]_*i*_ and [*Cl*^−^]_*o*_ are elevated to about **145 mM** to eliminate Goldman rectification [19]. According to our GNP framework, to ensure a strictly linear I-V relationship, [*Cl*^−^]_*i*_ should be 2 mM higher than [*Cl*^−^]_*o*_, given that 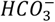 ions have asymmetric concentrations. Thus, we assume [*Cl*^−^]_*i*_ = **147 *mM*** and [*Cl*^−^]_*o*_ = 145 *mM*. Under similar conditions, experimental studies report the single-channel conductance of ***GABA***_***A***_ channels to be around **30 pS** [20, 39]. Based on these data, we can calculate out the permeabilities of ***GABA***_***A***_ channels for chloride ion 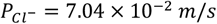 and for bicarbonate ion 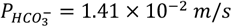 (refer to Eq. 16).

Under physiological conditions, [*Cl*^−^]_*i*_ is much lower than [*Cl*^−^]_*o*_, which creates outward rectification in ***GABA***_***A***_ channels and voltage-dependent conductance [39]. With the estimated single-channel permeability, we can reproduce the I-V curve of a single ***GABA***_***A***_ channel (Eq. 19) and determine the corresponding conductance (see Eq. 20 in Methods), as shown in Fig. 3a (diamond) and 3b (solid black curve). Notably, the conductance calculated using Eq. 20 is not equivalent to the slope of the I-V curve except when the voltage equals the channel’s reversal potential (Fig. 3b, dotted line; see also Supporting Information Fig. 1).

**Figure 3.**
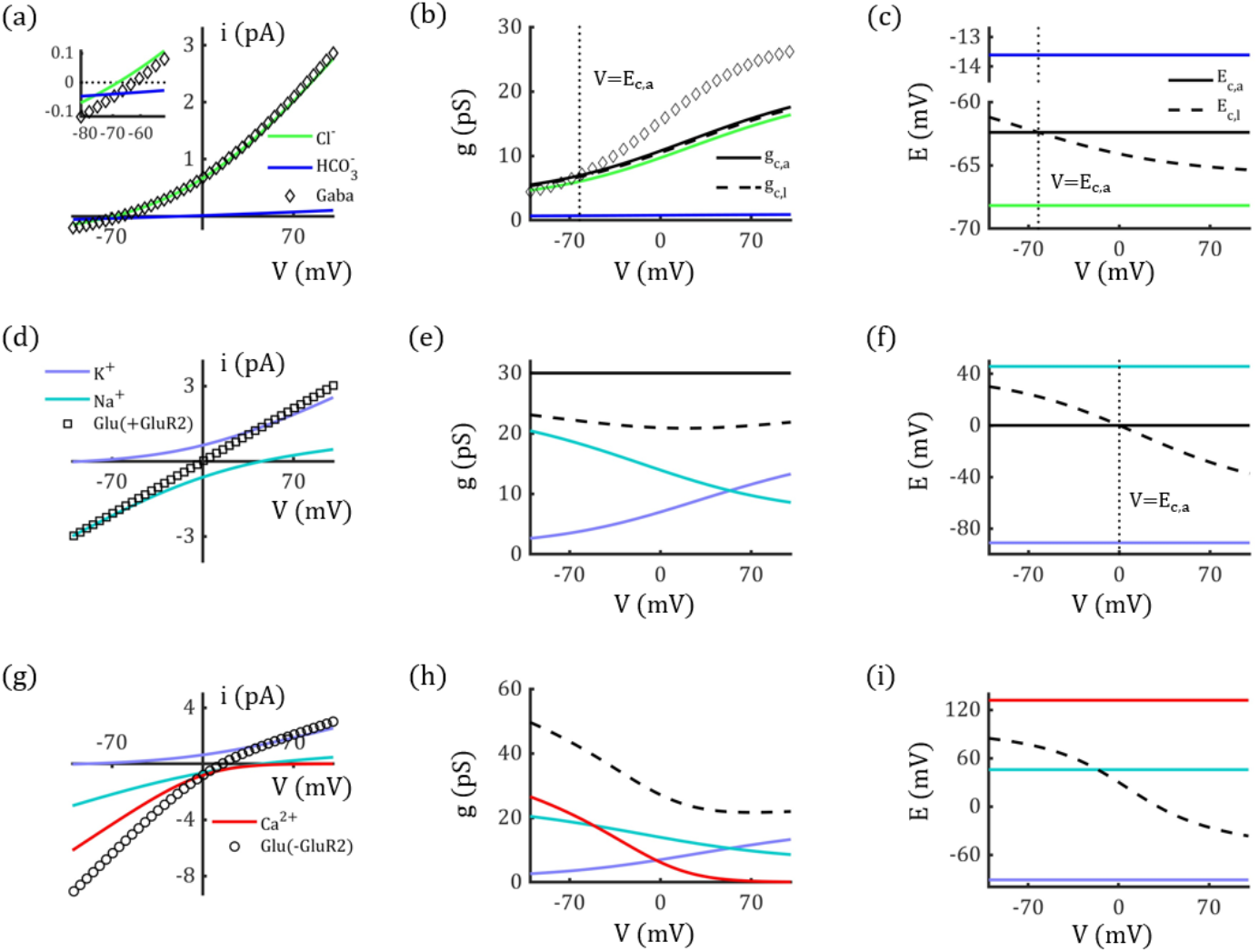
Rectification of *GABA*_*A*_ and *AMPA* Channels. (a-c) Outward rectification of single ***GABA***_***A***_ channel (a), corresponding conductance (b), and reversal potentials (c). Inset figure: curves are zoomed in around the reversal potential. (d-f) Linear I-V relation of single GluR2-containing ***AMPA*** channel which is impermeable to calcium ions, corresponding conductance (e), and reversal potentials (f). (g-i) Inward Rectification of single GluR2-lacking ***AMPA*** channel which is permeable to calcium ions, corresponding conductance (h), and reversal potentials (i). Here the calcium concentrations are set to be **[*Ca***^**+**^]_***o***_ **= 2 *mM*** and **[*Ca***^**+**^]_***i***_ **= 1 × 10**^**−4**^ ***mM***, and the permeability is 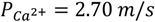. In (b, e): Two methods are used to calculate single channel conductance: *g*_*c,a*_ (solid black line) and *g*_*c,l*_ (dashed black line). In (c, f): Two methods are used to represent the channels’ reversal potentials: *E*_*c,a*_ (solid black line) and *E*_*c,l*_ (dashed black line). In (h,i): *g*_*c,a*_ and *E*_*c,a*_ are not applicable to the GluR2-lacking ***AMPA*** channel which is permeable to calcium ions, as calcium has different valence from potassium and sodium ions, whereas, *g*_*c,l*_ and *E*_*c,l*_ are still valid to describe the I-V curve (dashed black line). The diamond curve in (b) represents the slope of the channel’s I-V curve; the vertical dotted lines in (b), (c), and (f) represent *V* = *E*_*c,a*_. The I-V curves represent −*i*_*c*_ and −*i*_*q*_, respectively, to align with experimental formats.

Furthermore, using the permeability of each permeant, we can determine the chloride and bicarbonate currents and conductance, as shown in Fig. 3a and 3b (colored curves). Notably, the currents and conductance ratios are neither equal to the permeability ratio 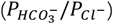 nor constant.

#### *AMPA* Channel

As another example, we investigated the properties of ***AMPA*** channels, which are permeable to both potassium and sodium ions. The corresponding concentrations are assumed to be **[*K***^**+**^]_***i***_ **= 96.83 *mM*, [*K***^**+**^]_***o***_ **= 3.17 *mM*, [*Na***^**+**^]_***i***_ **= 23.58 *mM*, [*Na***^**+**^]_***o***_ **= 131.42 *mM*** [36]. For these concentrations, the permeability ratio 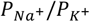 needs to be 0.87 in ***AMPA*** channels to ensure exact linear I-V relationship. Based on experimental data [32, 40], the single-channel conductance and pore diameter of ***AMPA*** channels are comparable to those of ***GABA***_***A***_ channels (**30 *pS*, 5 Å**). Using Eq. 16, the permeabilities of ***AMPA*** channels are estimated to be 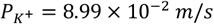 and 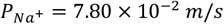.

We reproduced the linear I-V curve for an individual ***AMPA*** channel using the estimated permeabilities, as shown in Fig. 3d (square). ***AMPA*** channels are typically characterized by a linear I-V relationship and are generally considered non-rectifying. However, as shown in Fig. 3d, our results suggest that the apparent linearity of ***AMPA*** currents may result from a counterbalance between the outward rectification of potassium currents and the inward rectification of sodium currents. This implies that rectification would naturally arise in **AMPA** channels if the counterbalance is disrupted due to changes in ion concentrations or permeability ratio. The conductance of potassium and sodium currents are illustrated in Fig. 3e (colored curves). Again, the ratios of currents and conductance do not equal the ratio of permeability, nor are they constant.

***AMPA*** channels exhibit inward rectification when they have high calcium permeability due to the absence of the GluR2 subunit [33]. GluR2-lacking ***AMPA*** channels are highly permeable to calcium ions, leading to a calcium current that exhibits strong inward rectification. As shown in Fig. 3g, the emergence of a calcium component (with conductance comparable to potassium and sodium currents, see Fig. 3h) disrupts the balance between potassium and sodium currents, ultimately causing the channel to display inward rectification. While substantial evidence suggests that this rectification is primarily due to polyamine block, the contribution of calcium currents remains unclear. Further investigation is needed to fully understand this phenomenon, including studies in calcium-free conditions to isolate the influence of calcium currents.

Notably, the “apparent” conductance *g*_*c,a*_ of *Ca*^2+^-permeable ***AMPA*** channels (see Eq. 19 and black solid curves in Fig. 3b, e) cannot be calculated in the same way as for ***GABA***_***A***_ and GluR2-containing ***AMPA*** channels, since calcium ions have a different valence compared to sodium and potassium ions. Interestingly, an alternative conductance formulation (see Eq. 23) effectively characterizes these channels and remains valid for channels with different permeant valences. This “latent” conductance *g*_*c,l*_ is represented by black dashed curves in Fig. 3b, 3e, and 3h, while the corresponding latent reversal potentials *E*_*c,l*_ are shown as black dashed curves in Fig. 3c, 3f, and 3i.

Importantly, both latent conductance and reversal potential are voltage-dependent, even for channels with linear I-V relationships. The apparent conductance *g*_*c,a*_ and reversal potential *E*_*c,a*_ should always be paired with the latent conductance *g*_*c,l*_ and latent reversal potential *E*_*c,l*_ when describing channel properties. However, previous studies have often mismatched these parameters. A detailed discussion of apparent and latent conductance and reversal potentials is provided in the Methods and Supporting Information sections.

### Modeling Neural Dynamics by GNP Approach

The results presented in the previous sections were obtained under steady-state conditions (see Methods). To further incorporate the dynamics of electrodiffusive ion transport into neural firing dynamics, such as neural discharges, we assume that the electrodiffusive dynamics within ion channels are much faster than the voltage dynamics (see Quasi-Static Assumption in Supporting Information).

Thereafter, we build a neural dynamics model based on our GNP method, where, the electrodiffusive ion transport would induce ion concentration changes, then altering membrane potential. The changes of ion concentrations and membrane potential would in turn modulate the electrodiffusive dynamics, forming a closed feedback loop, as shown in Fig. 4. Our GNP approach explicitly incorporates electrodiffusive dynamics by modifying membrane conductance ***G***_***q***_, enabling a more comprehensive neurodynamic model that accounts for rectifying channels (Fig. 4b). Moreover, the GNP model captures the direct connection between ion concentration changes and neural activity, highlighting the role of passive-active ion transport interactions in neural stability. The simulation results and analysis by this model are displayed in following sections.

**Figure 4.**
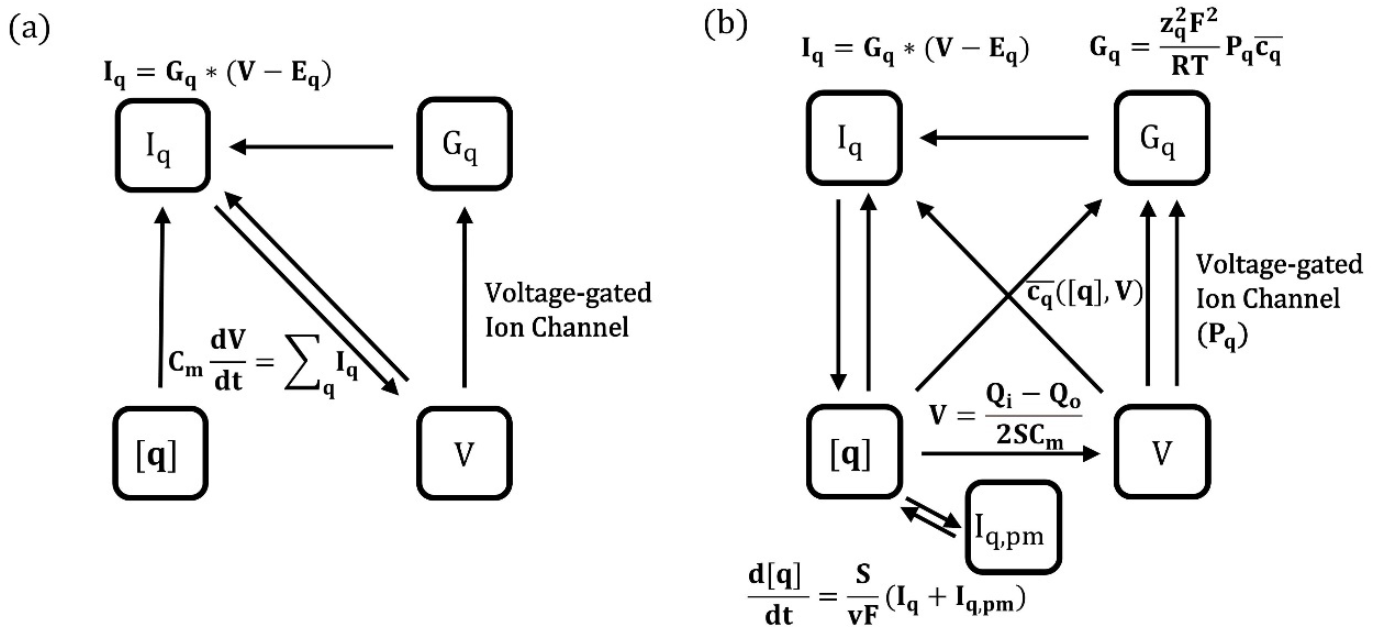
Comparison Between Classical Equivalent-Circuit-Based Model and Our GNP Model. Panels (a) and (b) depict the dynamic schematics of the classical equivalent-circuit-based model and our electrodiffusion-based GNP model, respectively. ***I***_***q***_: electrodiffusive ion current through ion channels; ***I***_***q***,***pm***_ : current induced by ***Na***^**+**^**/*K***^**+**^ **− *ATPase*** pumps; ***V*** : membrane potential; **[*q*]**: ion concentrations; ***G***_***q***_: membrane conductance per unit area; ***Q***_***i*/*o***_: net charge in intra- and extracellular spaces; 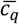: harmonic mean of intramembrane concentration profile.

#### Outwardly Rectifying Leaky Current Enhance Neural Stability

Leaky currents arise from various ion channels within the membrane [17, 41, 42]. In our model, we simplify these currents as an equivalent “leaky channel” characterized by high potassium permeability 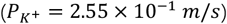, medium chloride permeability 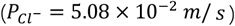, and low sodium permeability 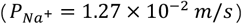. This leaky channel exhibits outward rectification, as shown in Fig. 5a-c.

**Figure 5.**
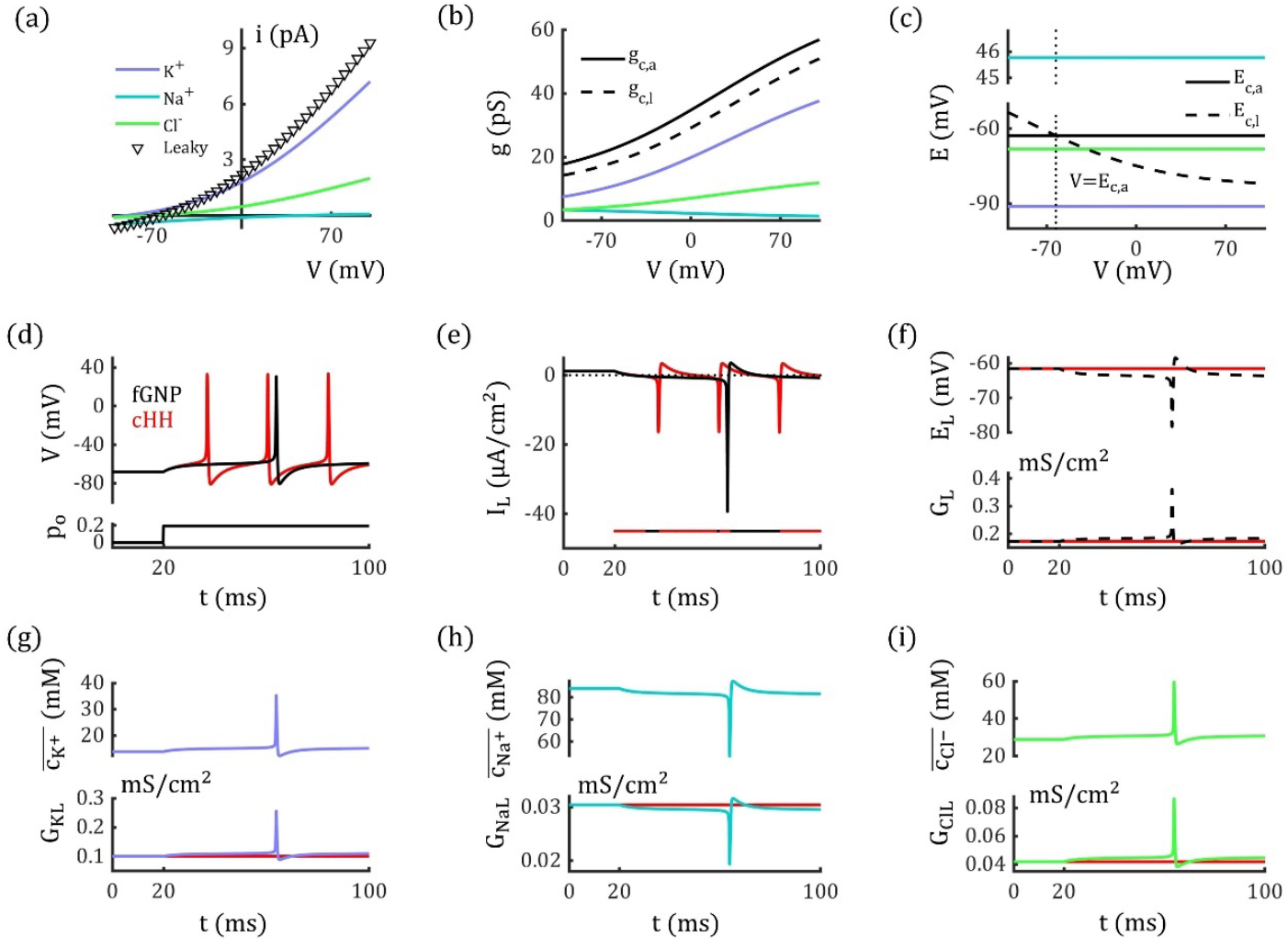
Outward Rectification of Leaky Channels Enhances Neural Stability. (a) The outward rectification of single leaky channel (triangle) used in fGNP model, where, each ion current is shown in purple (***K***^**+**^), cyan (***Na***^**+**^) and green (***Cl***^***−***^) curves. (b) Conductance of leaky channel as function of voltage. Solid black line represents ***G***_***c***,***a***_, dashed black line represents ***G***_***c***,***l***_, colored lines represent the conductance of respective ions. (c) Reversal potential of the leaky channel. Solid black line represents ***E***_***c***,***a***_, dashed black line represents ***E***_***c***,***l***_, colored lines represent reversal potentials of respective ions. (d) The neuron in the fGNP model generates fewer action potentials than the cHH model with same tonic glutamate stimuli. Black line represents concentration-fixed GNP (fGNP) model with rectifying leaky channel, red line represents the classical HH-type (cHH) model with linear leaky channel. (e) Leaky currents in the fGNP and cHH models during stimulation, with the horizontal line indicating the model with higher leaky current at each time point. The dotted line represents **0 *μA*/*cm***^**2**^. (f) Conductance of the leaky channel in the fGNP model. Dashed black lines represent ***G***_***c***,***l***_and ***E***_***c***,***l***_in fGNP model, red lines represent constant ***G***_***LH***_ and ***E***_***LH***_ in cHH model. (g-i) Harmonic mean and conductance of potassium and chloride ions increase, for sodium ions they decrease accompanying with the depolarization.

**Figure 6.**
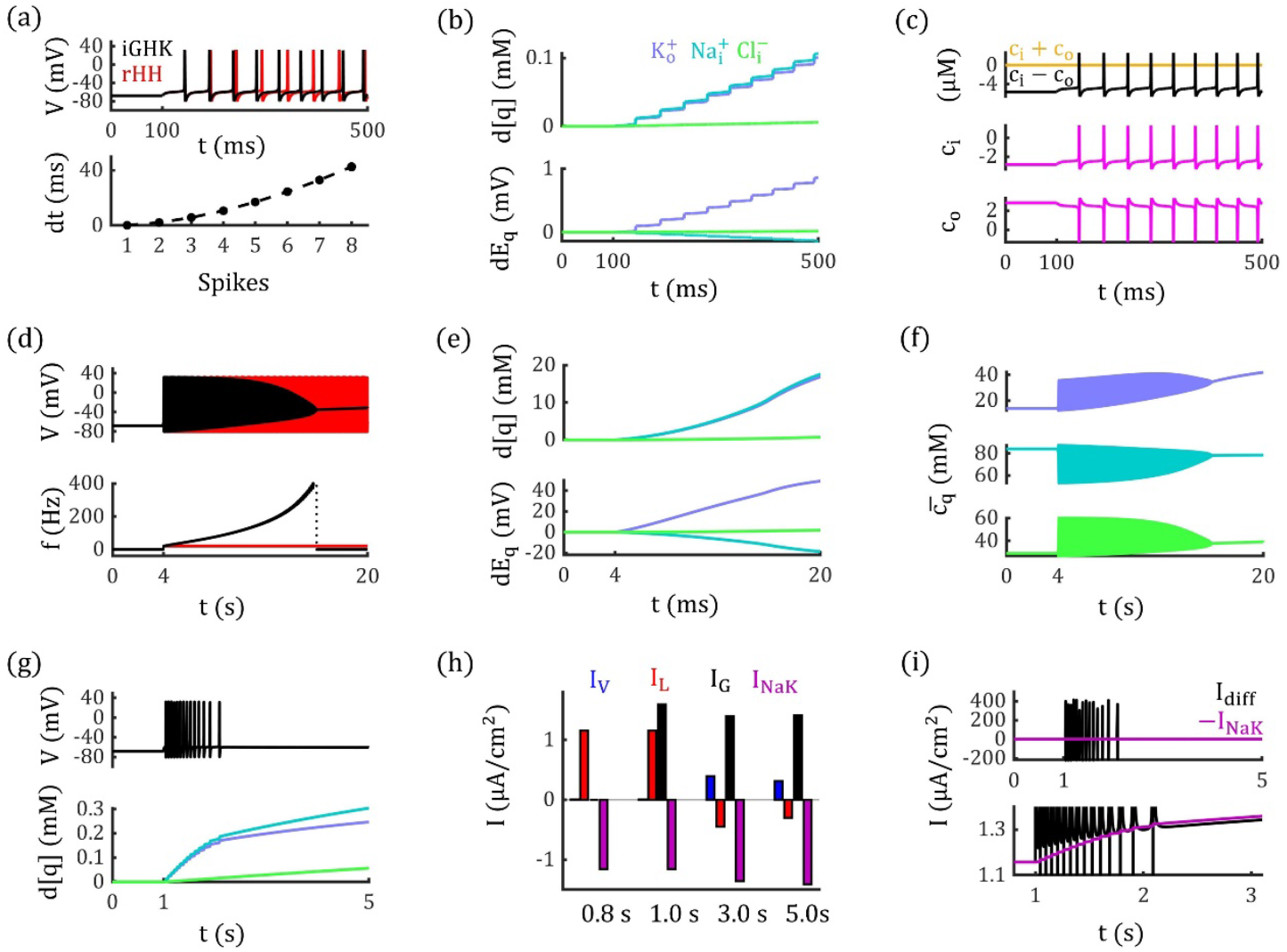
Differential Neural Dynamics in the GNP Model Incorporating electrodiffusion and Ion Concentration Changes. (a-f): Constant ***Na***^**+**^**/*K***^**+**^ **− *ATPase*** activity. (a) Upper panel: Neural discharge events in response to glutamate stimuli (***p***_***o***_ **= 0.20**) in the GNP (black) and fGNP (red) models. Lower panel: The timing of discharge events in the GNP model progressively advances the spikes in fGNP model. (b) The deviations of ion concentrations from their steady-state values gradually increase during neural activity (Upper panel), resulting in rising potassium and chloride equilibrium potentials and decreasing sodium equilibrium potential (Lower panel). (c) Dynamics of net charge concentrations in the intra- and extracellular spaces (***c***_***i***_ and ***c***_***o***_ magenta lines); their summation and difference are displayed in the upper panel. (d) In the GNP model (black), the firing frequency progressively increases, eventually leading to depolarization block under prolonged stimuli (***p***_***o***_ **= 0.20**). In contrast, the neuron in the fGNP model (red) maintains a constant firing frequency under the same stimuli. (e) Significant changes in ion concentrations and reversal potentials occur under prolonged stimulation, following the same trend as shown in (b). (f) The harmonic means of intramembrane potassium and chloride concentrations increase, whereas for sodium, it decreases during neural activity. (g-l): ion-concentration dependent ***Na***^**+**^**/*K***^**+**^ **− *ATPase*** activity. (g) With concentration-dependent ***Na***^**+**^**/*K***^**+**^ **− *ATPase*** activity, the neuron in the GNP model returns to a resting state after a few action potentials under glutamate stimulation (*p*_*_ = 0.20). (h) Currents from voltage-gated channels (***I***_***V***_), leaky channels (***I***_***L***_), glutamate channels (***I***_***G***_), and ***Na***^**+**^**/*K***^**+**^ **− *ATPase* (*I***_***NaK***_) at different time points. (i) The ***Na***^**+**^**/*K***^**+**^ **− *ATPase*** current (purple) increases, counterbalancing the electrodiffusive current (*I*_4)55_), thereby restoring neural stability.

Unlike the classical Hodgkin-Huxley (cHH) models [43], which assume constant conductance of leaky channel, our GNP model incorporates outwardly rectifying leaky currents. To explicitly see the effect of this rectification on neural dynamics, we fix ion concentrations in the GNP model to steady state (see Tab. 1) and compare the neural behavior in this concentration-fixed GNP (fGNP) model to that in the cHH model (see Concentration-Fixed GNP Model in Methods). To determine the average membrane permeability, we assume there are **1 × 10**^**7**^ leaky channels and **4 × 10**^**6**^ ***AMPA*** channels (see Fig. 3a) per square centimeter of membrane [9, 44], with an opening probability (*p*_*o*_) of ***AMPA*** channels to represent the intensity of glutamate stimuli. Voltage-gated currents are modeled using reduced Traub-Miles (RTM) equations (see Methods) [45].

As shown in Fig. 5d, the fGNP and cHH models respond differently to identical glutamate stimuli. With ***p***_***o***_ increasing from **0** to **0.20**, the neuron in the fGNP model fires only one action potential within **80 *ms***, while the neuron in the cHH model fires three. Due to rectification, the latent leaky conductance ***G***_***L***,***l***_ in the fGNP model rises to higher values than ***G***_***LH***_ in the HH model during depolarization (Fig. 5e). Specifically, potassium (Fig. 5g) and chloride (Fig. 5i) leaky conductance increases, while sodium conductance decreases (Fig. 5h). These changes cause ***G***_***L***,***l***_ to lower the leaky reversal potential ***E***_***L***,***l***_ (according to Eq. 25) in the fGNP model compared to ***E***_***LH***_ in the cHH model (Fig. 5e, 5f). As a result, the overall leaky current in the fGNP model is typically lower than in the cHH model (Fig. 5e, 5f), thus reduces neural excitability.

#### Electrodiffusion Promotes the Interplay between Ion Accumulation and Neural Discharge Events

When incorporating ion concentration dynamics, our GNP model uncovers that subtle ion concentration changes can evoke remarkable voltage signals. As illustrated in Fig. 6a, 6b, under tonic glutamate stimulation with ***p***_***o***_ **= 0.20**, ion concentrations in the GNP model are changed due to enhanced electrodiffusive ion transport, leading to a sequence of action potentials. Interestingly, despite significant variations in membrane potential, the corresponding ion concentration changes remain surprisingly small, with all changes below **1 %** after nine spikes (Fig. 6b).

Notably, the concentrations of net charges in intra- and extracellular spaces (***c***_***i***_ and ***c***_***o***_) are on the micromolar scale (Fig. 6c), which are much smaller than the millimolar scale of ion concentrations. As a result, even seemingly negligible ion concentration changes are sufficient to alter the charge difference significantly, thereby dramatically impact neural spike timing. As demonstrated in Fig. 6a, the timing of discharge events in the GNP model (black line) deviates significantly from the concentration-fixed model (red line) after just nine spikes. Such deviations could influence the neural temporal coding and impact on neural functions [46].

It is important to note that the ion concentration accumulation contributes to the increased neural excitability. As shown in Fig. 6a, under tonic glutamate stimulation (***p***_***o***_ **= 0.20**), the enhanced electrodiffusive ion transport gradually advances the timing of discharge events in the GNP model compared to the fGNP model. With prolonged glutamate stimulation of the same strength, these currents further elevate the firing frequency, ultimately drives the neuron into a depolarization block (DB) state (Fig. 6d black curve), while the identical stimuli elicit a constant firing activity in the fGNP model (Fig. 6d red curve).

Furthermore, before entering the DB state, both firing frequency and ion concentrations accelerate coherently in GNP model, indicating a positive feedback loop between ion accumulation and neural discharge events. Although changes in conductance and the sodium reversal potential tend to reduce excitability (Fig. 6e, 6f), this feedback loop is primarily driven by the elevation of potassium and chloride reversal potentials (Fig. 6e), which significantly weakens the inhibitory effects of potassium and chloride currents.

#### Interplay of Electrodiffusive and Active Ion Transports Determines Neurodynamic State

In Fig. 6d, the neuron generates high-frequency spiking and eventually enters the DB state. This transition occurs because the external glutamate stimuli disrupt the initial balance between passive ion transport by electrodiffusion and active ion transport by ***Na***^**+**^**/*K***^**+**^ **− *ATPase*** activity, which destroys the neural stability and lead to excessive excitability. To restore stability, the activity of ***Na***^**+**^**/*K***^**+**^ **− *ATPase*** needs to increase during neural activity to prevent excessive excitability and maintain normal function. Experimental studies have shown that ***Na***^**+**^**/*K***^**+**^ **− *ATPase*** activity is enhanced when extracellular potassium **[*K***^**+**^]_***o***_ and intracellular sodium **[*Na***^**+**^]_***i***_ levels rise [47–49]. To model this process, we implement concentration-dependent ***Na***^**+**^**/*K***^**+**^ **− *ATPase*** based on previous research (see Methods) [48, 49].

As illustrated in Fig. 6g, introducing concentration-dependent ***Na***^**+**^**/*K***^**+**^ **− *ATPase*** allows the neuron in the GNP model to regain a resting state after generating several action potentials by glutamate stimuli (***p***_***o***_ **= 0.20**). During this phasic-spiking event, ***Na***^**+**^**/*K***^**+**^ **− *ATPase*** activity increases by approximately **20 %**, with the pump-induced current rising from **−1.16 *μA*/*cm***^**2**^ at ***t* = 1 *s*** to **−1.41 *μA*/*cm***^**2**^ at ***t* = 5 *s*** (Fig. 6h). This increased ***Na***^**8**^**/*K***^**8**^ **− *ATPase*** activity counterbalance the electrodiffusive currents (Fig. 6i), eventually results in a resting state.

The response of ***Na***^**+**^**/*K***^**+**^ **− *ATPase*** activity to ion concentration changes influences neural dynamics under tonic glutamate stimulation. In addition to the phasic-spiking event shown in Fig. 6g, we identified three other types of neural behaviors arising from the interplay between passive and active ion transport. As shown in Fig. 7a, with relatively weak glutamate stimulation at ***p***_***o***_ **= 0.10**, the neuron undergoes slight depolarization. However, the depolarization amplitude gradually decreases following stimulus onset.

**Figure 7.**
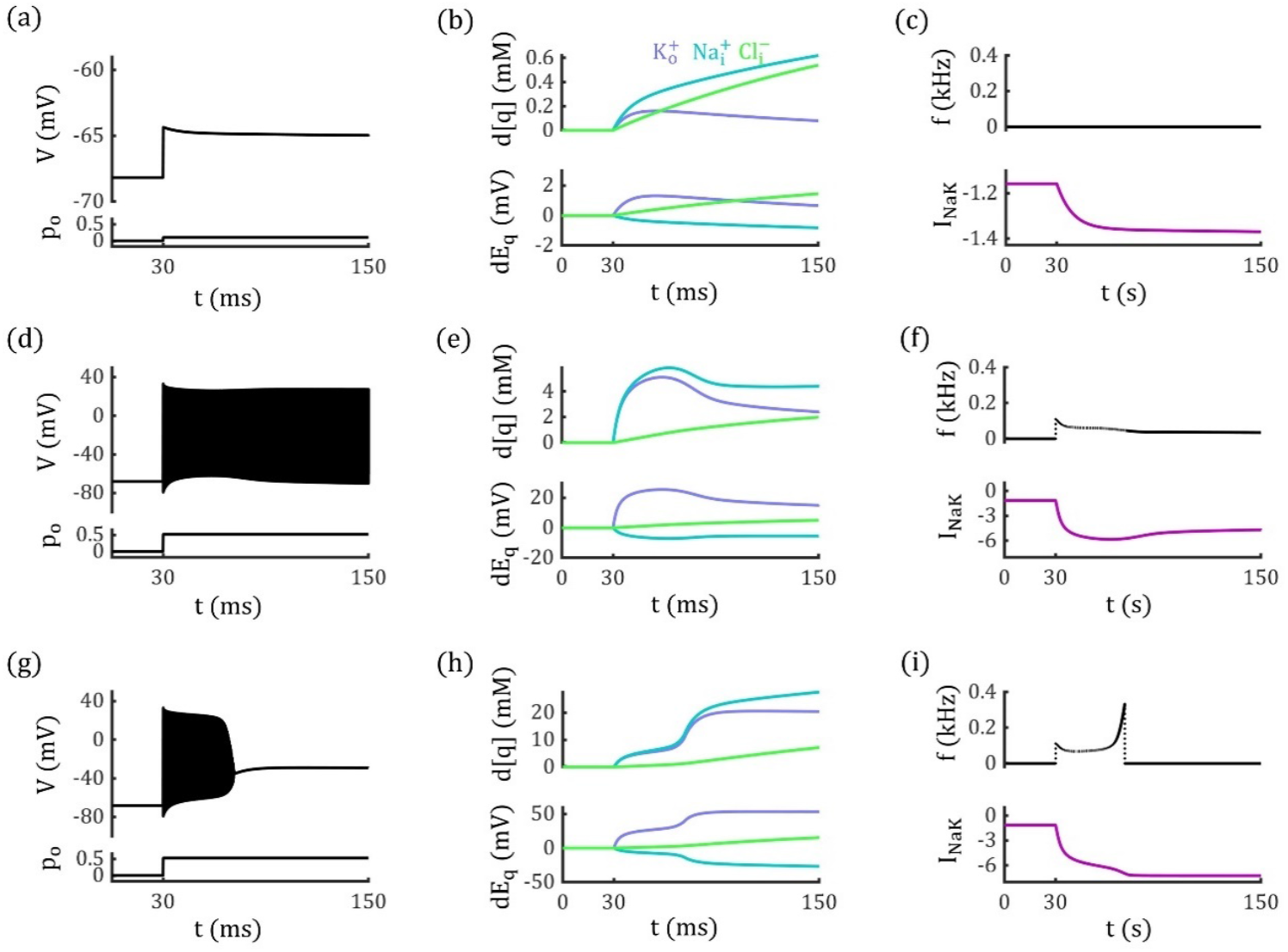
Competition Between Passive and Active Transport Settles Neural activities. (a-c) At ***p***_***o***_ **= 0.10**, the glutamate input enhances passive transport and induces depolarization, but the subsequent increase in ***Na***^**+**^**/*K***^**+**^**− *ATPase*** suppress the passive transport, maintaining resting state. (d-f) At ***p***_***o***_ **= 0.52**, the passive electrodiffusive current and active ***Na***^**+**^**/*K***^**+**^**− *ATPase*** activity reach a dynamic equilibrium, resulting in tonic spiking behavior. (g-i) At ***p***_***o***_ **= 0.53**, the electrodiffusive current overwhelms ***Na***^**+**^**/*K***^**+**^**− *ATPase*** activity, leading the neuron into a DB state. Left column: The glutamate input and the neural activity; Middle column: Changes in ion concentrations and corresponding reversal potentials. Right column: Neural firing frequencies and the changes in pump-induced currents.

With stronger glutamate stimuli at ***p***_***o***_ **= 0.52**, the ***Na***^**+**^**/*K***^**+**^ **− *ATPase*** activity and electrodiffusive currents achieve a dynamic equilibrium, resulting in a tonic-spiking state (Fig. 7d-7f). As shown in Fig. 6d, under ***p***_***o***_ **= 0.52**, neural discharge increases **[*K***^**+**^]_***o***_ and **[*Na***^**+**^]_***i***_, which in turn enhances the pump current ***I***_***NaK***_ (Fig. 7e, 7f). The elevated ***I***_***NaK***_ reduces the neural firing frequency (Fig. 7f) and cccgradually decreases **[*K***^**+**^]_***o***_ and **[*Na***^**+**^]_***i***_, counterbalancing the passive electrodiffusive currents and leading to a stable tonic-spiking state.

When the stimulus intensity is further increased to ***p***_***o***_ **= 0.53**, the electrodiffusive currents become too strong to be suppressed by the ***Na***^**+**^**/*K***^**+**^ **− *ATPase*** activity, eventually pushing the neuron into the DB state (Fig. 7g-7i). As shown in Fig. 7i, the firing frequency initially decreases due to the enhanced pump current ***I***_***NaK***_. However, the upper limit of pump strength makes ***I***_***NaK***_ fails to counterbalance the enhanced passive transport, and eventually leads the neuron to the DB state.

#### Electrodiffusive Dynamics Modulate the Voltage-gate Currents and Neural Dynamics

Traditionally, the dynamic equations governing the gating variables of voltage-gated channels are fit into conductance-based circuit equations (***I***_***qV***_) [43, 45], whereas gating equations for permeability-based GHK format remain underdeveloped. In above-mentioned GNP model, we simply employed the original RTM equations and ignored the electrodiffusive dynamics within voltage-gated channels. This approach is only valid under minor variations in ion concentrations, as the voltage-dependent conductance changes can be sufficiently captured by the gating equations. However, it may reduce accuracy when modeling neural activities with significant concentration changes, as the concentration-dependence of conductance is overlooked in original RTM equations.

To investigate the potential impact of electrodiffusive dynamics on voltage-gated currents and neural dynamics, we model the voltage-gated currents using the GHK format (***I′***_***qV***_) (see Methods). The I-V curves of ***I′***_***qV***_ in GHK format differ from those in the RTM model. To align them in steady state, we introduce a function ***f***_***q***_**(*V*)** to fit 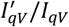, as illustrated in Fig. 8a, 8b. After modified by ***f***_***q***_**(*V*)**, the equivalent maximum conductance of the voltage-gated channels ***G′***_***KM***_ and ***G′***_***NaM***_ exhibits both voltage- and concentration-dependencies, as depicted in Fig. 8c.

**Figure 8.**
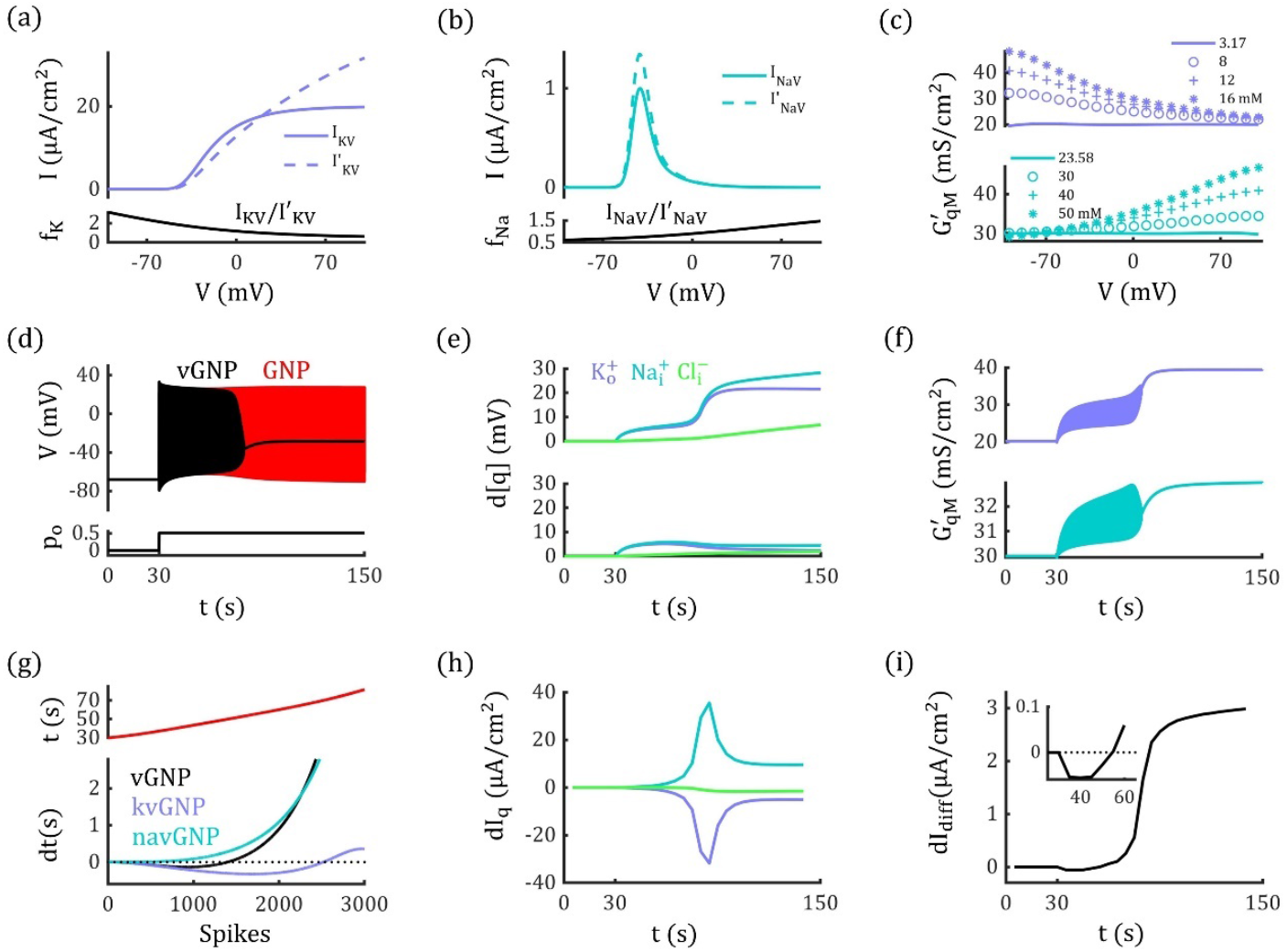
Electrodiffusive Voltage-Gated Currents Modulate Neural Activity. (a) and (b): Upper panel: I-V curves for voltage-gated currents in GHK format 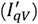 and RTM format (***I***_***qV***_) with ion concentrations fixed at steady-state levels. Lower panel: The ratios of ***I***_***qV***_**/*I′***_***qV***_, which are fitted by function ***f***_***qV***_**(*V*)** (see Methods). (c) Equivalent maximum conductance of voltage-gated potassium and sodium currents in the vGNP model. (d) Neuron enters the depolarization block state in the vGNP model (black line), while the neuron in the GNP model (red line) shows tonic-spiking by the identical glutamate stimuli (***p***_***o***_ **= 0.52**). ***Na***^**+**^**/*K***^**+**^**− *ATPase*** strength is concentration-dependent in both models. (e) Deviations in ion concentrations from their steady-state values in the vGNP model (upper panel) and GNP model (lower panel). (f) Maximum conductance of voltage-gated potassium and sodium currents both increases during the simulation in vGNP model. (g) Timing differences in neural discharge events in vGNP (black), kvGNP (purple), navGNP (cyan) models relative to the GNP model, illustrating the lag and lead in action potential generation. Upper panel: Timing of discharge events in the GNP model. (h) Differences of electrodiffusive potassium, sodium, and chloride currents between the vGNP and GNP models. (i) Differences of total electrodiffusive current between the vGNP and GNP models. In (a) and (b), the fitted parameter values are: **[*N***_***K*3**_, ***N***_***K*2**_, ***N***_***K*1**_, ***N***_***K*0**_**] = [−1.20 × 10**^**7**^, **1.18 × 10**^**−5**^, **0.0075, 0.85]** and **[*N***_***Na*3**_, ***N***_***Na*2**_, ***N***_***Na*1**_, ***N***_***Na*0**_**] = [9.41 × 10**^**−8**^, **4.64 × 10**^**−6**^, **−0.0056, 1.11]** (see Eq. 44). Data in (h) and (i) are processed using a sliding window (see Methods).

We incorporate ***I′***_***qV***_ into the GNP model and named it as the vGNP model, which shows different neural dynamics from the GNP model. As shown in Fig. 8d, with ***p***_***o***_ **= 0.52**, the neuron in the GNP model exhibits tonic-spiking, whereas the neuron in the vGNP model evolves into a DB state. This disparity is due to substantial changes in ion concentrations (Fig. 8e), which induce pronounced alterations in the conductance of ***G′***_***KM***_ and ***G′***_***NaM***_, as demonstrated in Fig. 8f.

In Fig. 8g, the neural discharge events in the vGNP model initially lag behind those in the GNP model because enhanced potassium conductance leads to stronger inhibition. However, by approximately 50 *s*, the discharge events in the vGNP model catch up with the GNP model as enhanced sodium conductance becomes dominant. This interplay between enhanced potassium and sodium conductance is further elucidated through the difference of each ion current (Fig. 8h) and total electrodiffusive currents (Fig. 8i).

Interestingly, in a kvGNP model (incorporating ***I′***_***KV***_ and ***I***_***NaV***_, see Methods), enhanced inhibitory potassium conductance can paradoxically increase neural excitability during certain periods (Fig. 8g). This occurs because the elevated conductance mitigates the inhibitory effects of potassium currents by promoting extracellular potassium accumulation and elevating 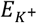. Similarly, in a navGNP model (incorporating ***I***_***KV***_ and ***I′***_***NaV***_), reduced sodium currents can also be observed in certain scenarios, as illustrated in Supporting Information Fig. 3.

## Discussion

This study investigates electrodiffusive dynamics of intramembrane ions on channel rectification and neural firing activities by an efficient Gauss-Nernst-Planck computational framework. We have clarified the relationship between conductivity and permeability with intramembrane ion concentration profiles, which makes critical step to bridge the gap between the permeability-based GHK equation and the conductance-based model. Meanwhile, our GNP method provides a practical approach to determine the permeabilities and conductance of ion channels to each permeant ions, demonstrated with ***GABA***_***A***_ And ***AMPA*** channels, which reveals that channel rectification arises from the superimposition of individual permeant ion currents. Our results demonstrate that electrodiffusive ion transport leads to rectifying leaky currents and enhances neural stability. We emphasize that the typically overlooked electrodiffusive dynamics can significantly influence neural activity through a feedback loop involving electrodiffusive ion transport, ion concentration changes, and membrane potential, critical in our model but absent in classical models. We also found that the competition between electrodiffusive ion transport and ***Na***^**+**^**/*K***^**+**^ **− *ATPase*** activity plays a crucial role in determining neurodynamic states by excitatory stimuli, such as resting, tonic-spiking, and DB states.

Our GNP framework provides promising potential for broad applications. One area of interest is the role of cellular impermeable anions in maintaining concentration homeostasis and neural functions. Traditionally, these anions have been considered important for osmotic pressure regulation [7, 50], but their impact on neural discharge events remains largely unexplored. Some studies suggest that local impermeable anions may play a role in establishing chloride concentrations, which are crucial for the inhibitory effects of ***GABA*** transmitters [51]. However, this conclusion remains a topic with intensive debate. Traditional conductance-based models are insufficient for exploring this problem, as impermeable anions do not generate transmembrane currents. In contrast, our GNP model incorporates impermeable anion concentrations as a critical factor in determining membrane potential (see Eq. 27), providing an appropriate framework to investigate how variations in these anions affect steady-state conditions and modulate neural activity.

Importantly, our model is well-suited for studying neural dynamics during epileptic seizures, which involve complex discharge patterns such as periodic bursting and DB, accompanied with significant ion concentration oscillations [36]. These dynamic features are challenging for traditional conductance-based models to capture, but our GNP framework—integrating concentration-dependent membrane conductance and the competition between passive and active transport processes—offers a powerful scheme for modeling the neural dynamics during epileptic seizures.

Our GNP framework has certain limitations, one of which is the exclusion of complex biophysical mechanisms such as phosphorylation [52], polyamine block [33, 34], and environmental interactions [26, 30], which could have cooperated with electrodiffusive ion transport to modulate channel properties. For example, we demonstrated that incorporating electrodiffusive dynamics within voltage-gated channels in the vGNP model led to altered neural activity compared to the standard GNP model (Fig. 8), indicating that voltage-gated channel dynamics are more complex than previously understood. Therefore, we acknowledge that our modeling of voltage-gated currents is incomplete, but the intricate gating dynamics of these ion channels require experimental validation to fully understand the mechanism.

Another simplification in our model is regarding the dynamics of chloride ions. We omit the contributions of cation-chloride cotransporters such as ***KCC*2** and ***NKCC*1** [53, 54], as well as GABAergic stimuli, which leaves the leaky chloride current as the sole mechanism modulating chloride concentrations. Consequently, intracellular chloride accumulation is slower than the changes in extracellular potassium and intracellular sodium concentrations during neural activity (Fig. 6). However, the regulation of chloride concentrations plays a crucial role in regulating the inhibitory effects of GABAergic signaling [53, 54], which is critical for maintaining proper neural function. Even though this simplification reduces the model’s complexity, it provides a concise and general framework that can be applied and extended to specific topics. A more comprehensive version of the model, incorporating the chloride cotransporters and GABAergic stimuli, will be explored in future studies to investigate the mechanism of chloride dynamics in regulating neural physiology.

## Methods

### The Gauss-Nernst-Planck Approach

To model electrodiffusion, a common approach is to solve the Poisson-Nernst-Planck (PNP) equation, expressed as:

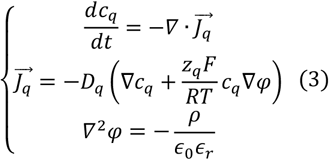

Here, ***c***_***q***_, ***φ*** and ***ρ*** represent the spatial distributions of ion concentration, electric potential, and charge density, respectively. The vector field 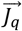 denotes the ion flux, while ***D***_***q***_ is the diffusion coefficient. The term ***ϵ***_**0**_***ϵ***_***r***_ represents the permittivity. The subscript ***q*** refers to ion species that permeate the neural membrane.

Solving the PNP equations is computationally demanding. Consequently, neural membranes are often approximated using an equivalent circuit model, described by the following equation:

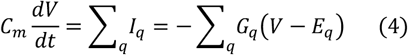

Equation (4) serves as a cornerstone for constructing practical neurodynamic models. Despite its success, the approximate equivalence between Eqs. (3) and (4) and the integration of electrodiffusive dynamics into conductance-based models remain open questions.

In the steady state, we find that for neural systems illustrated in Fig. 1, enforcing overall electrical neutrality and replacing Poisson’s equation with Gauss’s law simplifies the PNP problem to a one-dimensional form:

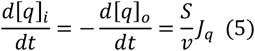

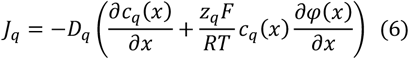

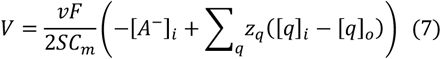

where ***q* = *K***^**+**^, ***Na***^**+**^, ***Cl***^**−**^. The concentration of impermeable anions, **[*A***^**−**^]_***i***_, is set to **110 *mM*** in this work. The full derivations are detailed in Supporting Information.

We analyze neural dynamics in a closed system, meaning the total intracellular and extracellular ion concentrations satisfy **[*q*]**_***i***_ **+ [*q*]**_***o***_ **= *constant***. Consequently, from Eq. 7, the membrane potential ***V*** evolves according to:

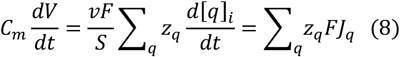

where ***J***_***q***_ is given by Eq. 6, which can be rewritten as:

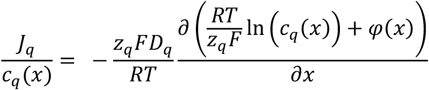

By integrating this equation from **0** to ***d***, we can obtain:

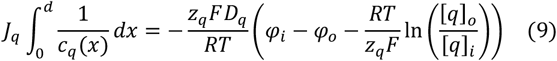

Combining Eqs. 8 and 9, we obtain:

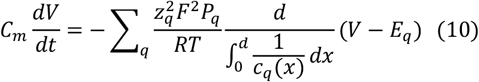

where ***P***_***q***_ **= *D***_***q***_**/*d*** defines the membrane permeability. From Eq. 10, it follows that the membrane conductance per unit area can be expressed as:

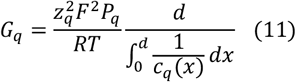

Eq. 11 reveals that membrane conductance ***G***_***q***_ is directly proportional to both permeability and the harmonic mean of the intramembrane concentration, which is defined as:

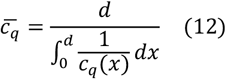

Given that the intramembrane concentration profile ***c***_***q***_**(*x*)** at steady state is obtained in the classical GHK approach [5, 6], which is:

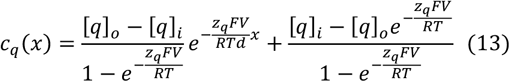

Using Eq. 13, we compute the harmonic mean concentration 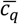 as Eq. 2:

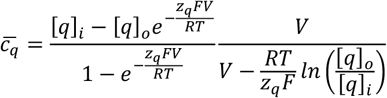

Notably, by combining Eqs 2, 11, and the circuit equation, we obtain the Goldman-Hodgkin-Katz current equation:

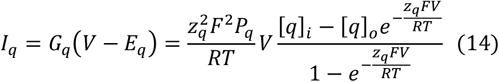

This demonstrates that our approach effectively bridges the gap between traditional cable models and the electrodiffusive framework. The Gauss-Nernst-Planck formulation provides a means to characterize the conductance of rectifying ion channels and seamlessly incorporate electrodiffusive dynamics into neurodynamic models.

### Characterizing channel rectifications

#### Condition for Linear I-V Relation

Due to electrodiffusive dynamics, single-channel openings may exhibit nonlinear I-V relationships. Many of these behaviors can be explained by the GHK equation, but quantifying the varying conductance remains challenging. Furthermore, it has been unclear how to separate the respective ionic currents when a channel is permeable to multiple ion species, as in ***GABA***_***A***_ and ***AMPA*** channels.

Our GNP approach provides a systematic way to address these issues. Consider ***GABA***_***A***_ And ***AMPA*** channels as examples. Both channels are permeable to two ion species. Due to electrodiffusive effects, these channels exhibit a linear I-V relationship under the following condition:

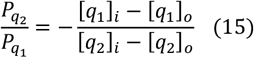

where ***q***_**1**_ and ***q***_**2**_ denote the two permeant ions, and 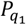 and 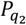 represent their respective permeabilities within a single open channel.

#### Channel Conductance and Permeabilities to Specific Permeants

***GABA***_***A***_ channels typically exhibit outward rectification. However, when measuring single-channel conductance, the intracellular and extracellular chloride concentrations are often adjusted to eliminate passive rectification. In contrast, ***AMPA*** channels generally display an approximately linear I-V relationship under physiological conditions. Once the linear I-V relationships of ***GABA***_***A***_ and ***AMPA*** channels are measured, the single-channel conductance ***g***_***c***,***m***_ can be determined from the slope of the I-V curve.

With ***g***_***c***,***m***_ known, the single-channel permeabilities can be computed using the following equations:

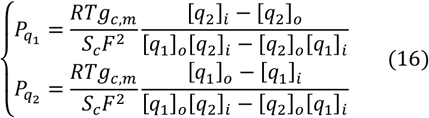

where ***S***_***c***_ represents the cross-sectional area of the channel pore. Experimental studies have rarely reported single-channel permeability data, yet permeability provides a more intrinsic characterization of channel properties compared to conductance, which depends on ion concentrations and membrane potential.

The individual conductance for ***q***_**1**_ and ***q***_**2**_ can be further derived as:

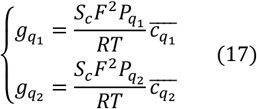

where 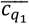 and 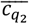 represent the harmonic mean concentrations of the respective ions.

Notably, the conductance ratio between ***q***_**1**_ and ***q***_**2**_ is given by:

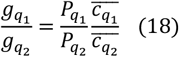

This result highlights that the conductance ratio does not necessarily equal the permeability ratio 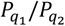, even though previous studies have often assumed them to be identical.

#### Conductance and Reversal Potential of Ion Channels

For an entire ion channel, the total current can be expressed as:

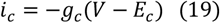

Interestingly, we found that conductance and reversal potential can be expressed in two distinct forms, both of which effectively characterize the channel’s electrical properties. We refer to the first form as the “apparent” conductance and reversal potential, given by Eqs. 20 and 21:

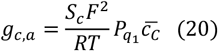

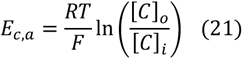

Here, ***q***_**1**_ represents an arbitrary permeant ion, and the intracellular and extracellular concentrations **[*C*]**_***i***_ and **[*C*]**_***o***_ are defined as:

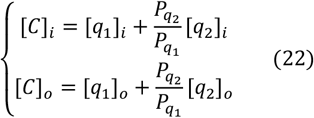

The term 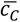 represents the harmonic mean of **[*C*]**, given by:

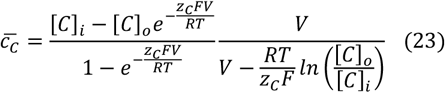

Here, ***z***_***C***_ is set to 1 for ***AMPA*** channels and **−1** for ***GABA***_***A***_ channels.

The apparent conductance ***g***_***c***,***a***_ and reversal potential ***E***_***c***,***a***_ remain constant under linear I-V conditions (see Eq. 15). However, when channels exhibit rectification, ***E***_***c***,***a***_ remains voltage-independent, whereas ***g***_***c***,***a***_ varies with membrane potential.

We have also identified another set of conductance and reversal potential, which we refer to as the “latent” conductance and reversal potential:

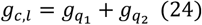

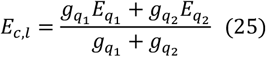

Unlike their apparent counterparts, both ***g***_***c***,***l***_ and ***E***_***c***,***l***_ are voltage-dependent, even when the I-V relationship is linear (see Fig. 3).

If we consider the entire membrane as a single effective “channel,” Eqs. 21 and 25 can be used to determine the resting membrane potential, corresponding to the Goldman-Hodgkin-Katz voltage equation and the chord conductance equation, respectively.

Crucially, apparent and latent conductance and reversal potentials must be carefully matched in pairs to accurately describe channel electrophysiological properties. However, previous studies have frequently mismatched these parameters, potentially leading to inaccurate interpretations and misleading conclusions. Detailed derivations and more generalized forms of Eqs. 15–25 are provided in the Supporting Information.

**Table 1.**
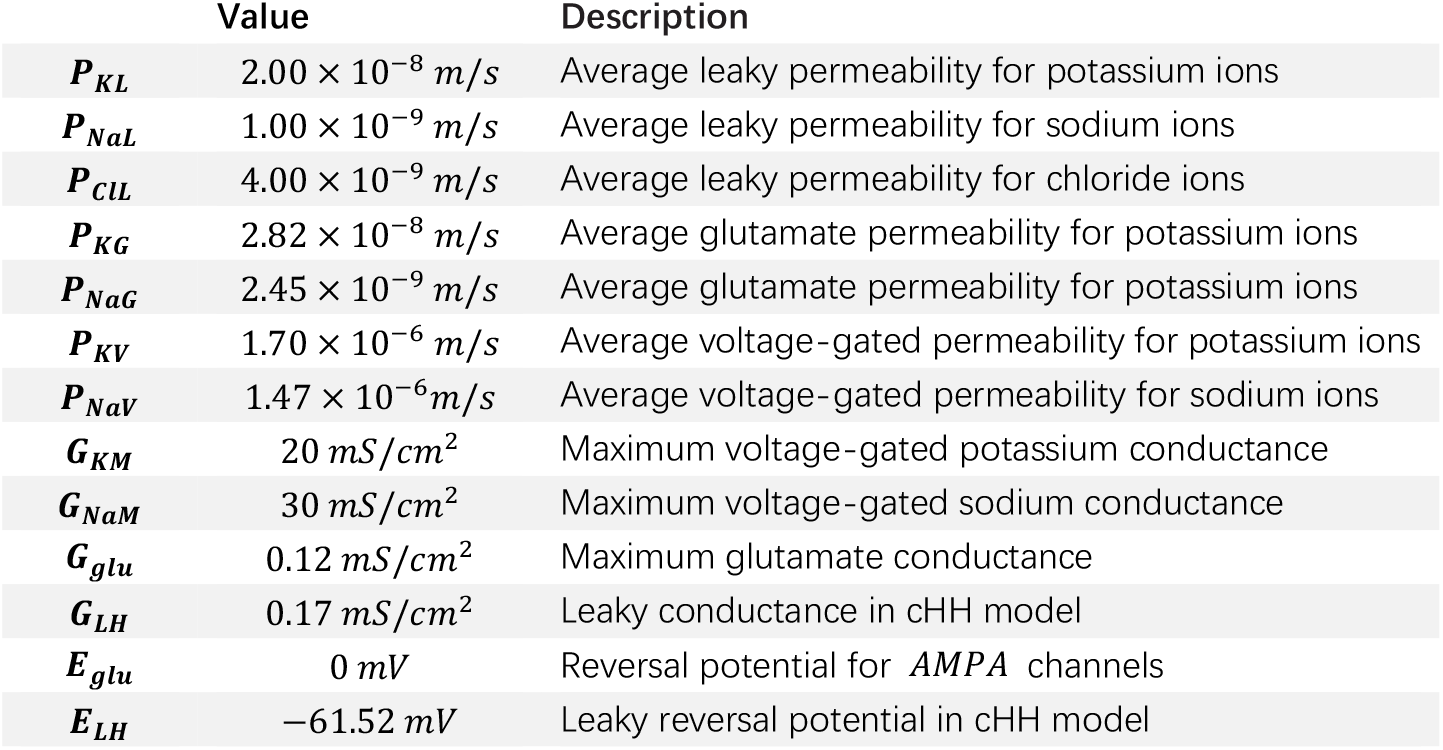
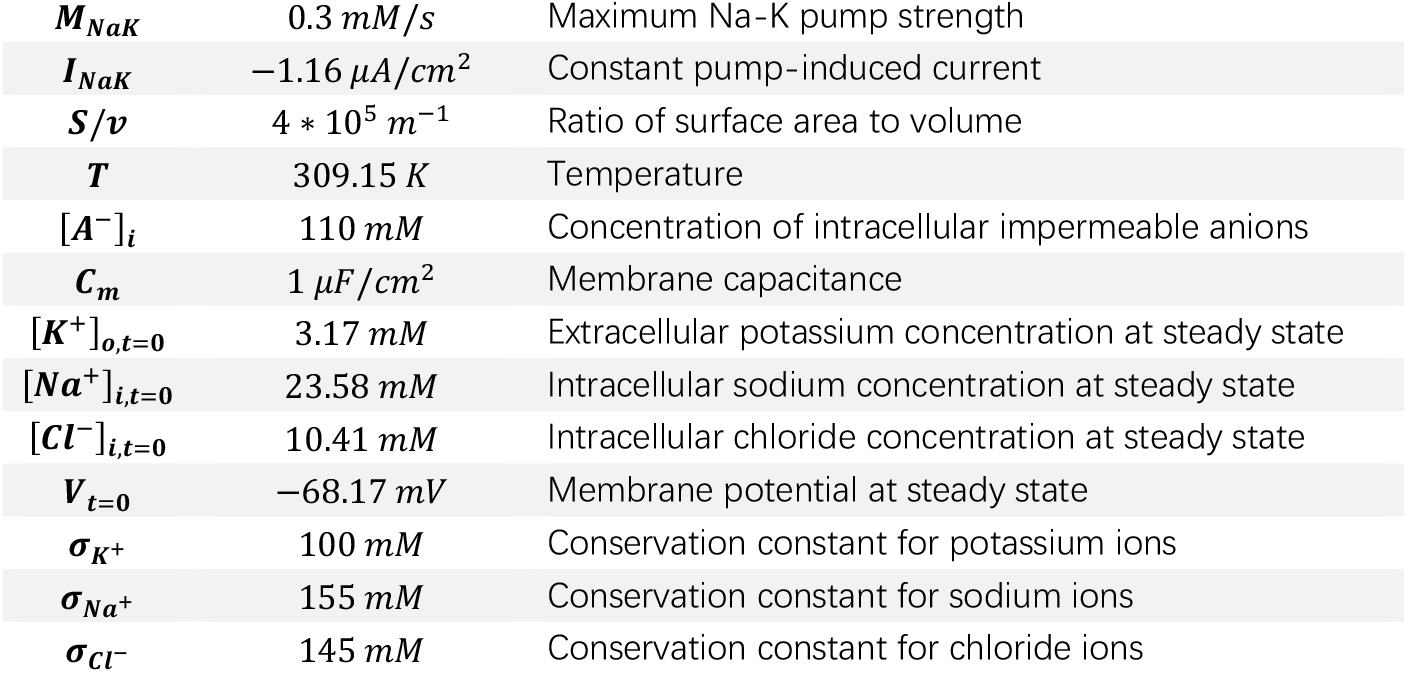
Units and description of the parameters used in the neurodynamic models.

### The Electrodiffusive Neurodynamic Model

#### The GNP Neurodynamic Model

Although the previous derivations and analyses are based on steady-state conditions, it is reasonable to extend these results to model neural dynamics, as intramembrane electrodiffusion occurs much faster than voltage dynamics (see Quasi-static Approximation in the Supporting Information). This allows us to incorporate channel rectifications into practical electrodiffusive neurodynamic models.

Our model includes three types of ion channels: voltage-gated channels, leak channels, and glutamatergic channels. Additionally, to balance passive ion transport and maintain ionic homeostasis, we incorporate active ion transport via the ***Na***^**+**^**/*K***^**+**^ **− *ATPase*** pump. These channels and pumps generate four types of ionic currents: voltage-gated current (***I***_***qV***_), leak current (***I***_***qL***_), glutamatergic current (***I***_***qG***_), and pump-induced current (***I***_***NaK***_).

Since the total concentration of each ion type remains constant in our isolated system (**[*q*]**_***i***_ **+ [*q*]**_***o***_ **= *constant***), we select extracellular potassium concentration (**[*K***^**+**^]_***o***_), intracellular sodium concentration (**[*Na***^**+**^]_***i***_), and intracellular chloride concentration (**[*Cl***^**−**^]_***i***_) as independent dynamic variables. The corresponding dynamic equations in our model are given by:

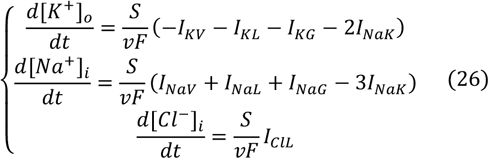

The **[*K***^**+**^]_***i***_, **[*Na***^**+**^]_***o***_, **[*Cl***^**−**^]_***o***_, and membrane potential ***V*** are determined by Eq. 27:

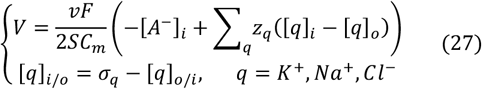

where ***σ***_***q***_ are constant parameters.

Leak channel currents are modeled using the GHK current equation:

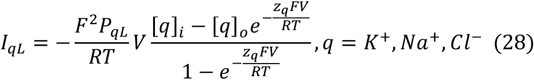

Similarly, ***AMPA*** channels currents are given by:

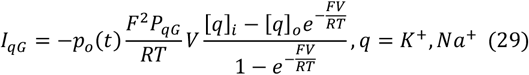

The ***P***_***qL***_ and ***P***_***qG***_ are averaged membrane permeabilities, which are calculated as:

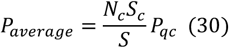

Here, ***P***_***qc***_ represents the single-channel permeabilities, estimated using Eq. 16. ***N***_***c***_**/*S*** denotes the number of channel *c* per unit area. The values of ***P***_***qc***_ and ***N***_***c***_**/*S*** are provided in the Results section, while the corresponding values of ***P***_***qL***_ and ***P***_***qG***_ are listed in Tab 1.

The corresponding conductance ***G***_***qL***_ and ***G***_***qG***_ is calculated using Eq. 2.

To investigate the interaction between passive and active ion transport, the pump-induced current ***I***_***NaK***_ is either held constant or varies with ion concentrations, as described by Eq. 31:

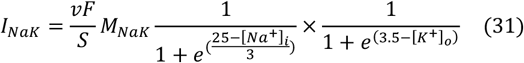

The voltage-gated currents are modeled as follows:

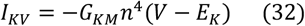

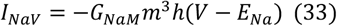

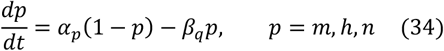

To model voltage-gated currents, we use a circuit-based approach and adopt the reduced Traub-Miles (RTM) equations for gating kinetics [45], as specialized gating kinetics for electrodiffusive models have not yet been developed. The gating variables *m, h, n* follow the RTM kinetics:

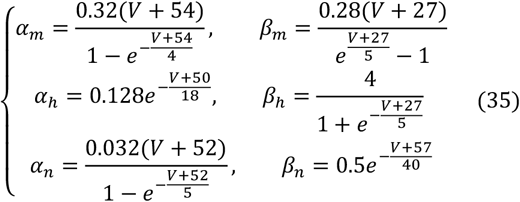

#### Concentration-Fixed GNP (fGNP) Model

Ion concentrations dynamically change during neural activity in this GNP neurodynamic model. However, for short-term neural activity, ion concentration changes may be negligible. In such cases, it is practical to fix ion concentrations and model neural activity using the following dynamic equation:

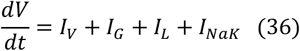

where the total voltage-gated current ***I***_***V***_, glutamatergic current ***I***_***G***_, and leak current ***I***_***L***_ are defined as:

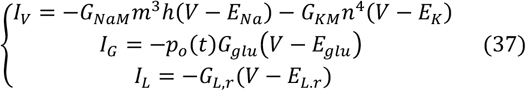

Here, ***G***_***NaM***_, ***G***_***KM***_, ***m, h, n*** share the same values and dynamic equations as described in Eqs. 32–35. Glutamate channels are assumed to exhibit a linear I-V relationship (see Eq. 15), with ***G***_***glu***_ and ***E***_***glu***_ calculated using Eqs. 20–23 (see Tab. 1). Leak channels, however, are assumed to exhibit rectifications due to the significantly higher membrane permeability to potassium than sodium. The latent voltage-dependent conductance ***G***_***L***,***r***_ and reversal potential ***E***_***L***,***r***_ are expressed as

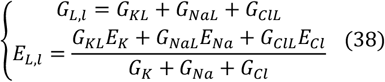

The specific leak conductance for potassium, sodium, and chloride is given by:

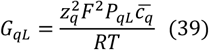

We define the total electrodiffusive current as the sum of voltage-gated, glutamatergic, and leak currents:

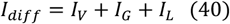

In Eq. 36, the active transport current ***I***_***NaK***_ is set as a constant. The model described by Eqs. 36–39 is referred to as the ion concentration-fixed GNP (fGNP) model.

The fGNP model closely resembles traditional conductance-based models, with the primary distinction being the introduction of rectifying leak channels (Eq. 39). To illustrate the impact of this modification, we also construct a classical Hodgkin-Huxley (cHH) model incorporating a linear leak current:

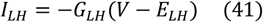

where ***G***_***LH***_ and ***E***_***LH***_ are constants, matched to ***G***_***L***,***l***_ and ***E***_***L***,***l***_ in the fGNP model at steady state. The currents ***I***_***V***_ and ***I***_***G***_ are identical in both the fGNP and cHH models.

In both the fGNP and GNP models, electrodiffusive effects within voltage-gated channels are neglected, an assumption valid only when ion concentration changes are minimal. However, when ion concentration dynamics become significant, this simplification becomes insufficient, as traditional RTM equations overlook the concentration dependence of conductance. To address this, we model voltage-gated currents using the GHK current equation:

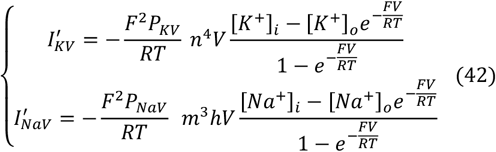

The gating variables ***m, h***, and ***n*** follow the same dynamics as in Eq. 35. Assuming the RTM equations accurately capture voltage-gated current rectification under steady-state ion concentrations, we introduce a normalization factor ***f***_***q***_**(*V*)** such that 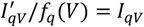 (at steady state). The function ***f***_***q***_**(*V*)** is fitted using a cubic equation (Fig. 8a, 8b, lower panels):

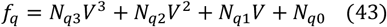

Using ***f***_***q***_**(*V*)**, the dynamic equations for extracellular potassium and intracellular sodium concentrations are:

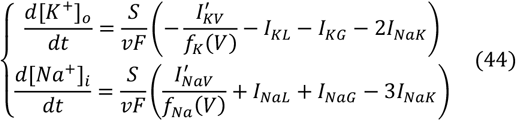

The model incorporating Eq. 44 is referred to as the vGNP model. The equivalent maximum conductance for voltage-gated currents in the vGNP model is given by:

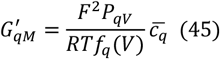

To analyze how variations in 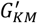 and 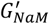 influence electrodiffusive currents, we compute the differences in ionic currents between the vGNP and GNP models:

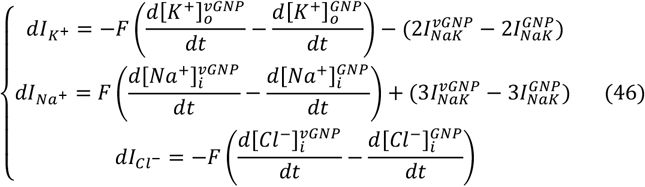

To smooth oscillations in the raw data, we apply a sliding window with an 8-second width and a 4-second step, as shown in Fig. 8h. The total difference in electrodiffusive currents, *dI*_*diff*_, is the sum of 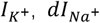 and 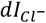, depicted in Fig. 8i.

To separately analyze the effects of 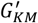 and 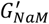on neural dynamics, we introduce two specialized models: kvGNP and navGNP.

kvGNP Model: Only the voltage-gated potassium current follows the GHK equation, while sodium currents are modeled using the RTM framework:

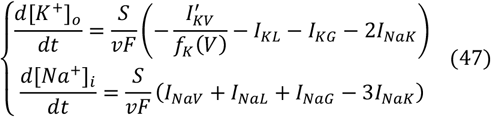

navGNP Model: Here, the sodium current follows the GHK equation, while potassium currents adhere to the RTM model

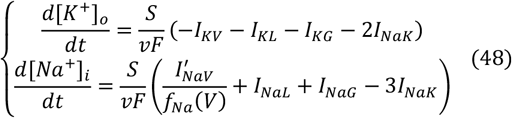

In both models, ***I***_***qV***_ follows the RTM model (Eq. 35). The dynamics of kvGNP and navGNP are illustrated in Supporting Information Fig. 3.

## Acknowledgement

We thank Prof. Wenxu Wang, Prof. Erik De Schutter, Dr. Jules Lallouette and Dr. Prabal Negi for helpful discussions. Zichao Liu is supported by Science and Technology Innovation 2030 major projects (No. 2021ZD0203803).

Yinyun Li is supported by Okinawa Institute of Science and Technology Graduate University.

## Supporting Information

### Establish Gauss-Nernst-Planck Model

Applying the overall electroneutrality assumption and Gauss’s law significantly simplifies the Poisson-Nernst-Planck problem in neural systems. Below, we outline how these conditions simplify the PNP equation (Eq. 3).

The total charge in the intracellular and extracellular spaces is given by:

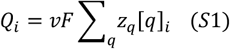

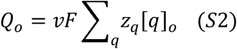

where ***q*** denotes a specific ion species with valence ***z***_***q***_ and concentrations **[*q*]**_***i***_ and **[*q*]**_***o***_ in the intracellular and extracellular spaces, respectively. ***F*** is Faraday’s constant, and *v* represents volume, which is assumed to be equal in both compartments.

To maintain overall electroneutrality, we require:

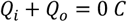

According to Gauss’s law, the electric field in the intramembrane space remains constant and is given by:

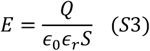

where ***Q* =** |***Q***_***i***_ **− *Q***_***o***_|**/2**, *S* is the surface area of the neural membrane within the intramembrane space (e.g., surface ***S*2** in Fig. 1a, 1b), and ***ϵ***_**0**_***ϵ***_***r***_ represents membrane permittivity. This formulation aligns with the constant-field theory [5, 6]. Crucially, using Gauss’s law allows us to avoid solving the Poisson equation directly. Instead, we can express the membrane potential as a function of ion concentrations:

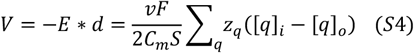

where ***d*** is the intramembrane distance, and ***C***_***m***_ **=** *ϵ*_**0**_***ϵ***_***r***_**/*d*** represents the membrane capacitance per unit area.

We assume the neural membrane is permeable to potassium, sodium, and chloride ions. Under physiological conditions, their typical intra- and extracellular concentrations are:

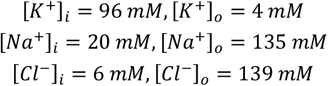

From these values, the intracellular and extracellular charge densities are:

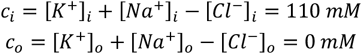

Substituting these into Eq. S4 results in an unrealistically high membrane potential exceeding **10**^**6**^ ***mV***, which is not observed in biological systems. This discrepancy highlights the necessity of including impermeable intracellular anions (***A***^**−**^) to balance the charge distribution. These anions are abundant in the intracellular space but scarce extracellularly. Thus, we introduce:

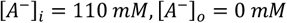

to maintain charge neutrality and produce a physiologically realistic membrane potential.

Notably, Eq. S4 establishes a direct relationship between voltage dynamics and ion concentration dynamics:

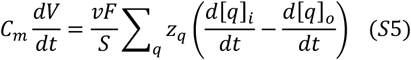

As illustrated in Fig. 1c, we assume ions are uniformly distributed within the intra- and extracellular spaces while continuously varying within the intramembrane space. At equilibrium, the diffusive flux ***J***_***qdiff***_ of ion *q* must remain constant across the neural membrane, as defined in Eq. S6:

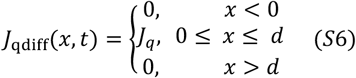

Under these conditions, the dynamic equations for **[*q*]**_***i***_ and **[*q*]**_***o***_ are:

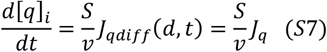

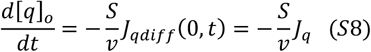

where ***J***_***q***_ is given by the Nernst-Planck equation (Eq. 6 in Methods).

These derivations, summarized in Eqs. 5–6 in Methods, form the foundation for constructing our electrodiffusive neurodynamic model.

### Characterizing Channel Properties

In experimental studies, the measured channel current (***i***_***c***_) and conductance (***g***_***c***_) are related to their respective current (***I***_***c***_, Eq. 18) and conductance (***G***_***c***_, Eq. 1) per unit area as follows:

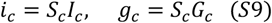

where ***S***_***c***_ represents the cross-sectional area of the channel’s pore domain.

For ion channels permeable to multiple ion types, the total channel current can be expressed as:

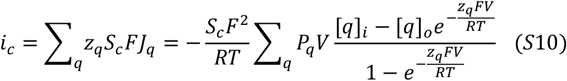

For channels permeable only to monovalent ions, Eq. S10 simplifies to:

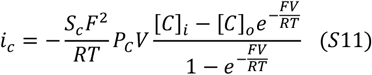

where ***P***_***C***_**[*C*]**_***i***_ and ***P***_***C***_**[*C*]**_***o***_ are constants. The value of ***P***_***C***_ is set equal to one of the permeant ion permeabilities (e.g.,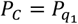). The equivalent concentrations **[*C*]**_***i***_ and **[*C*]**_***o***_ are defined as:

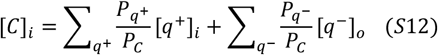

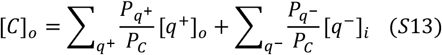

where ***q***^**+**^ and ***q***^**−**^ represent monovalent cations and anions, respectively.

When **[*C*]**_***i***_ **= [*C*]**_***o***_, the channel exhibits a linear I-V relationship, and Eq. S11 reduces to:

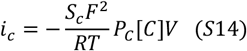

where **[*C*] = [*C*]**_***i***_. The apparent channel conductance, corresponding to the slope of the linear I-V curve, is given by:

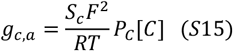

Equations S11–S15 provide a framework for estimating ion channel permeabilities from experimental data.

Consider a channel with two types of permeants, ***q***_**1**_ and ***q***_**2**_, both with the same valence. Setting 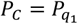, the ion concentrations and permeability ratio must satisfy Eq. S16 to achieve **[*C*]**_***i***_ **= [*C*]**_***o***_ in order to maintain a linear I-V relationship:

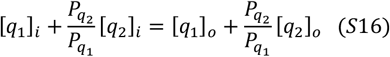

Solving for the permeability ratio:

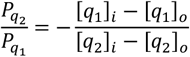

The total current then becomes:

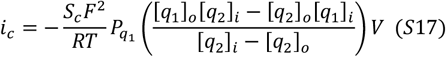

If ion concentrations **[*q*]**, single-channel conductance ***g***_***c***,***m***_, and pore diameter ***S***_***c***_ are known experimentally, the permeabilities 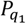 and 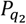 can be computed using Eq. 16:

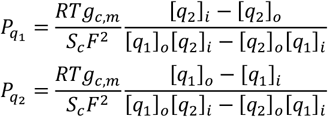

Once permeabilities are determined, the conductance for ***q***_**1**_ and ***q***_**2**_ can be calculated as Eq. 17:

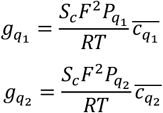

When **[*C*]**_***i***_ **≠ [*C*]**_***o***_, the channel exhibits rectification, and the total current is given by:

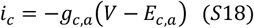

where the apparent conductance is:

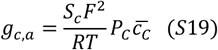

and the apparent reversal potential is:

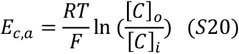

Interestingly, Eq. S18 can be rewritten in a circuit-like form:

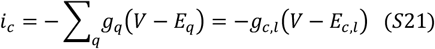

where we define the latent conductance and reversal potential as:

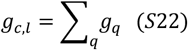

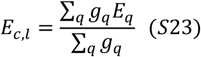

Both **(*g***_***c***,***a***_, ***E***_***c***,***a***_), and **(*g***_***c***,***l***_, ***E***_***c***,***l***_) can reproduce the channel’s I-V curve. However, while ***g***_***c***,***a***_ and ***E***_***c***,***a***_ are only valid for channels with equivalent permeants, ***g***_***c***,***l***_ and ***E***_***c***,***l***_ have no such limitation. Further details on apparent and latent conductance and reversal potentials are provided in Supporting Information Fig. 1.

**Supporting Information Figure 1.**
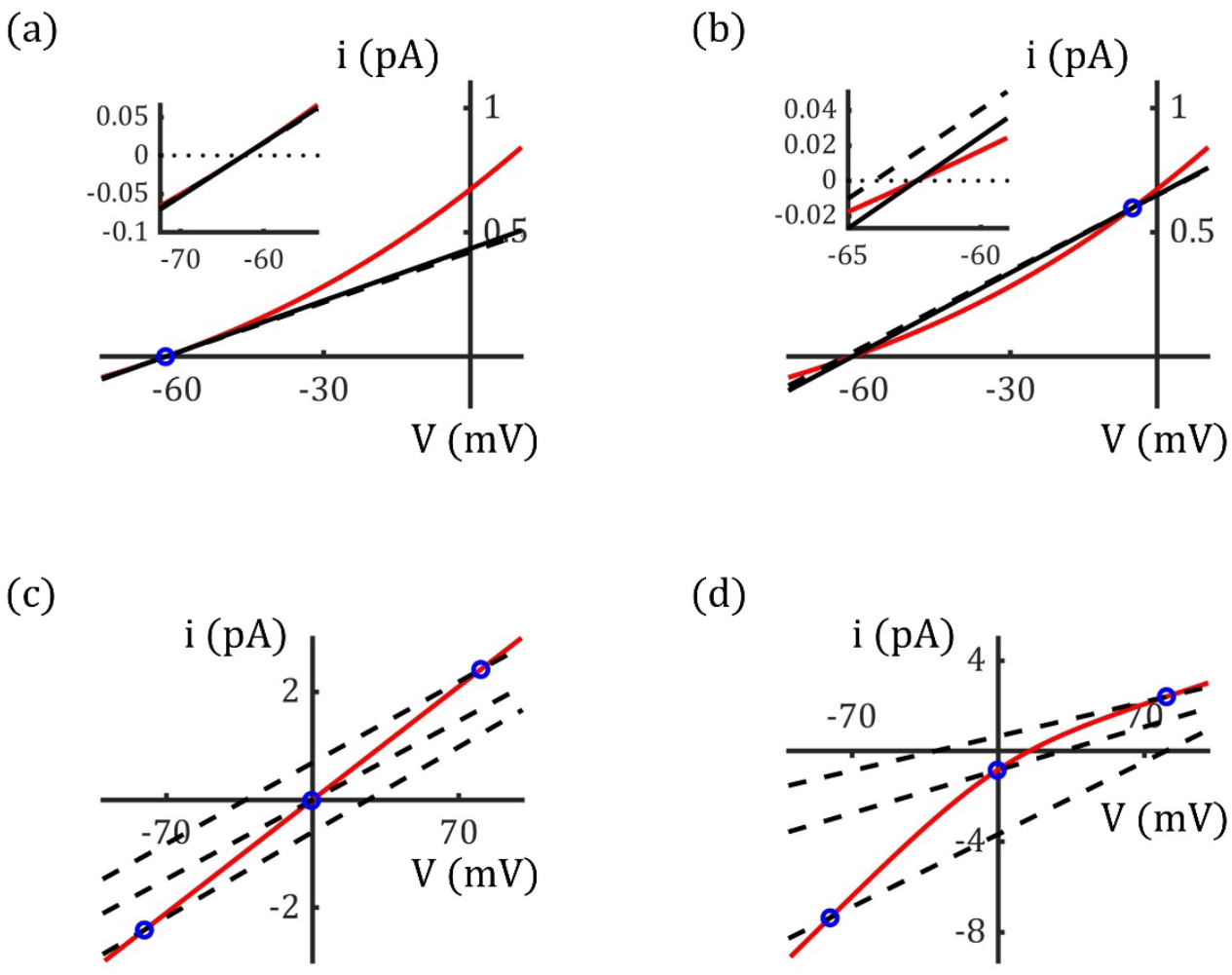
Apparent and Latent Conductance and Reversal Potentials Reproduce the I-V Curves of Ion Channels. (a) I-V curve for ***GABA***_***A***_ channel. At ***V* = −62.34 *mV***, the current through the ***GABA***_*V*_ channel is **0 *pA***, marked by the blue circle. The apparent and latent reversal potentials are identical and equal to ***V* = −62.34 *mV***, as shown at the inset. However, the apparent and latent conductance, ***G***_***r***,***a***_ and ***G***_***r***,***c***_, differs at this voltage, represented by the slopes of the black solid and dashed lines, respectively. The red curve depicts the current through the ***GABA***_***A***_ channel. (b) At ***V* = −5.00 *mV***, both the apparent and latent conductance increases. The apparent reversal potential remains constant, while the latent reversal potential (dashed black line) shifts downward, as shown at the inset. (c) The linear I-V relationship of an individual calcium-impermeable ***AMPA*** channel (red curve) can be characterized by the latent conductance and reversal potential, demonstrated here by three cases (black dashed lines) at ***V* = −80 *mV, V* = 0 *mV*** and ***V* = 80 *mV***. (d) An individual GluR2-lacking ***AMPA*** channel, permeable to calcium ions, exhibits inward rectification (red curve). The apparent conductance and reversal potential are not applicable in this case, but the latent conductance and reversal potential successfully reproduce the I-V curve, as illustrated by three cases (black dashed lines) at ***V* = −80 *mV, V* = 0 *mV*** and ***V* = 80 *mV***.

### Quasi-static Approximation

To investigate the impact of electrodiffusion on neural dynamics during transient neural activity, we assume that electrodiffusive dynamics within channel pores occur much faster than voltage dynamics. Consequently, these dynamics can be approximated as a series of “quasi-static” states during neural spikes. To validate this assumption, we estimate the characteristic timescale for the intramembrane concentration profile to transition from ***c***_***q***_**(*x, V***_**0**_) to ***c***_***q***_**(*x, V***_**0**_ **+ Δ*V*)** in response to a small membrane potential perturbation Δ*V*.

In general, the temporal evolution of the concentration profile within an ion channel follows:

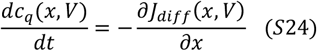

Assuming a steady state at ***V* = *V***_**0**_, we obtain:

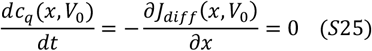

which implies to:

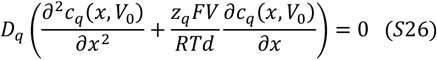

If the membrane potential shifts to ***V***_**0**_ **+ Δ*V*** while the concentration profile remains temporarily at ***c***_***q***_**(*x, V***_**0**_), the rate of change in concentration, ***dc***_***q***_**(*x, V*)/*dt***, can be approximated as:

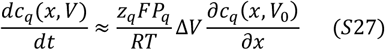

Meanwhile, the new steady-state profile ***c***_***q***_**(*x, V***_**0;**_ **+ Δ*V*)** can be expressed as:

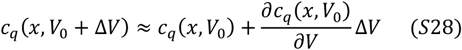

The characteristic time required for the concentration profile to shift from ***c***_***q***_**(*x, V***_**0**_) to ***c***_***q***_**(*x, V***_**0**_ **+ Δ*V***), denoted as **Δ*t***, can be estimated as:

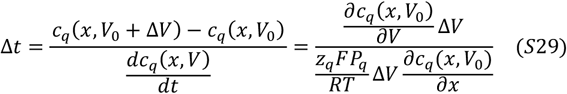

Reorganizing Eq. S29 gives:

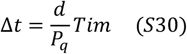

where ***Tim*** is expressed as:

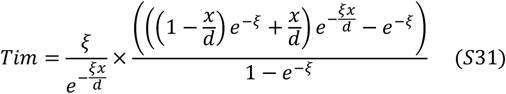

with 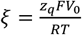. The function *Tim* reaches its maximum value at ***x* = *d*/2** when ***V***_**0**_ approaches **0 *mV***. Notably, for ***z***_***q***_ **= 1**, the maximum ***Tim*** is approximately 0.125 (see Supporting Information Fig. 2).

According to Eq. S30, **Δ*t*** is directly proportional to the membrane thickness ***d*** and inversely proportional to permeability ***P***_***q***_. As estimated in the main text, the permeabilities of ***GABA***_***A***_ and ***AMPA*** channels are on the order of **10**^**−2**^ ***m*/*s***. Given a membrane thickness of approximately **10 *nm*, Δ*t*** is estimated to be less than **10**^**−6**^ ***s***. Since typical membrane potential changes occur on a timescale of **10**^**−3**^ ***s***, this **Δ*t*** is negligible. This analysis supports the assumption that electrodiffusive dynamics can be approximated as a series of quasi-static states during dynamic neural events, justifying the application of steady-state equations (Eqs. 5–14) from our GNP model to describe neural dynamics.

**Supporting Information Figure 2.**
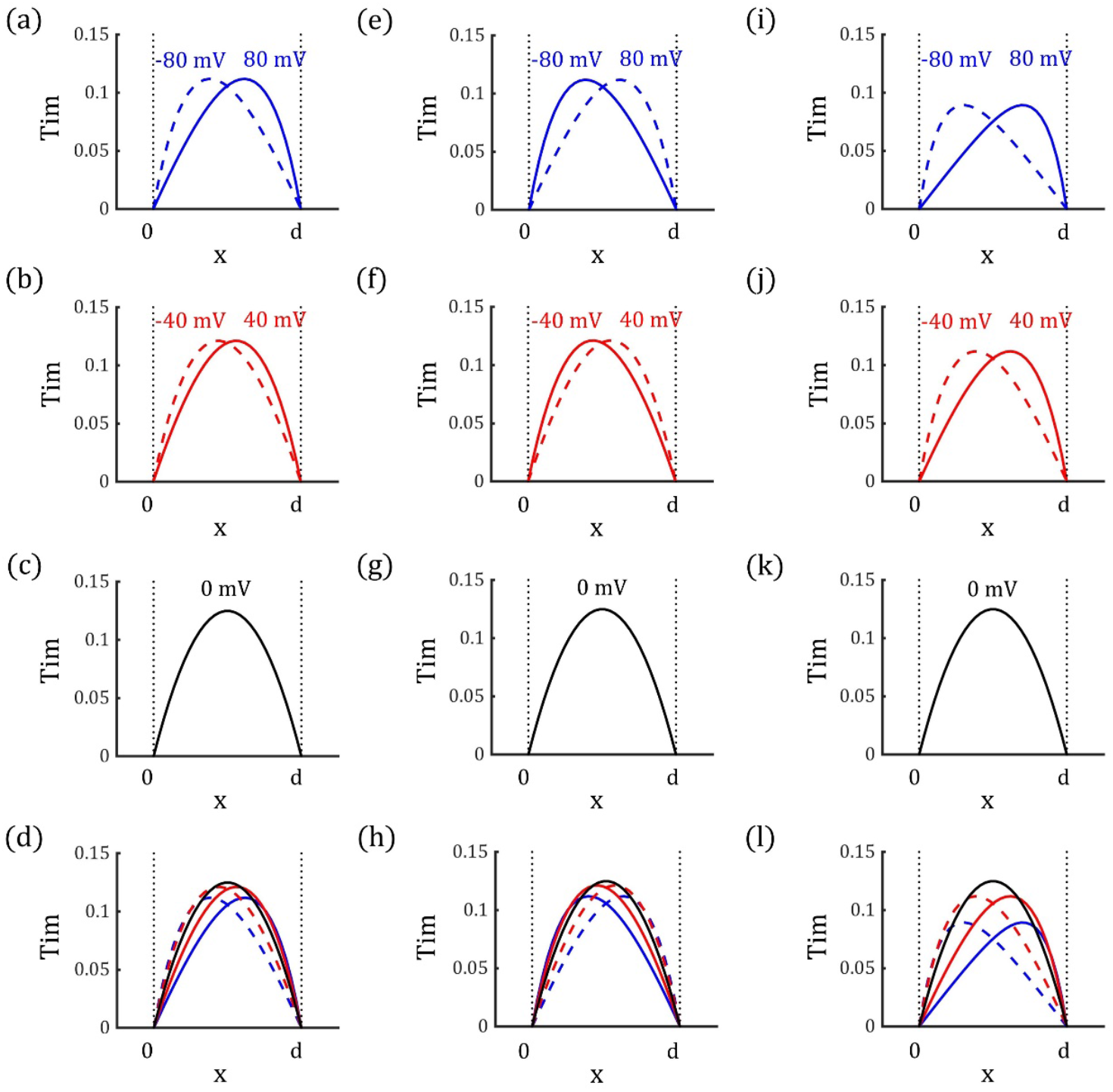
Values of *Tim* across Different Locations and Voltages. (a-d): For ***z***_***q***_ **= +1**. (e-h): For ***z***_***q***_ **= −1**. (i-l): For ***z***_***q***_ **= +2**. The function ***Tim*** reaches its maximum value of approximately 0.125 when ***V* = 0 *mV*** and ***x* = *d*/2.**

**Supporting Information Figure 3.**
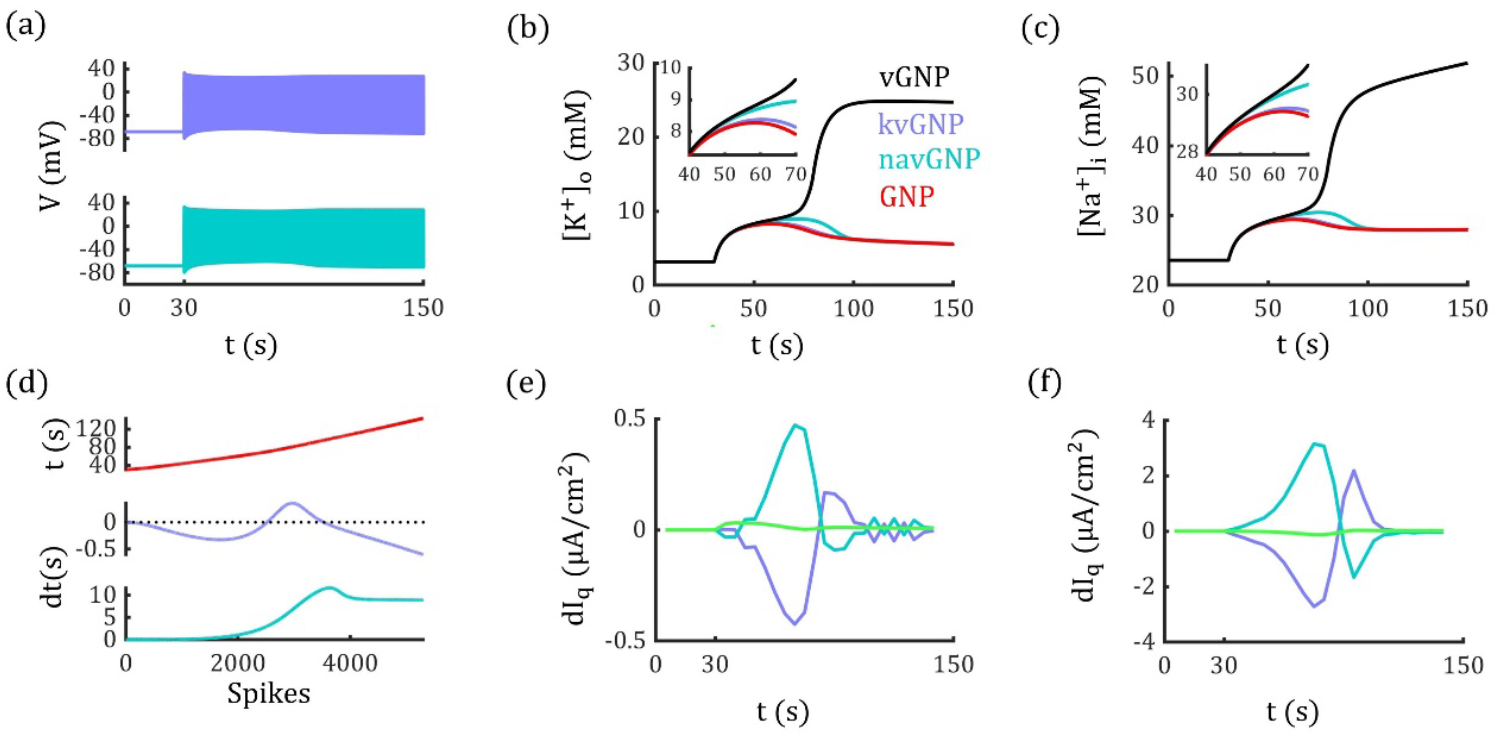
Neural Dynamics in kvGNP and navGNP Models in Response to Glutamate Stimuli. (a) Neural discharge events in kvGNP (upper panel) and navGNP (lower panel) models. The glutamate stimuli are identical to those used in Fig. 8d (***p***_***o***_ **= 0.52**). Both models exhibit tonic-spiking rather than entering a DB state. (b, c) Changes in extracellular potassium (b) and intracellular sodium (c) concentrations for four different models. Insets: zoomed in version of concentration changes between **40 *s*** and **70 *s***. (d) Timing differences in neural discharge events in kvGNP (middle panel), navGNP (lower panel) models relative to the GNP model, illustrating the lag and lead in action potential timing. Upper panel: Timing of discharge events in the GNP model. (e, f) Differences in electrodiffusive potassium, sodium, and chloride currents in kvGNP (e) and navGNP (f) models, relative to the GNP model. Note that in the kvGNP model, the potassium current can be less inhibitory than in the GNP model, while in the navGNP model, the sodium current may be less excitatory during certain periods.

